# Inferring allele-specific copy number aberrations and tumor phylogeography from spatially resolved transcriptomics

**DOI:** 10.1101/2024.03.09.584244

**Authors:** Cong Ma, Metin Balaban, Jingxian Liu, Siqi Chen, Li Ding, Benjamin J. Raphael

## Abstract

A key challenge in cancer research is to reconstruct the somatic evolution within a tumor over time and across space. Spatially resolved transcriptomics (SRT) measures gene expression at thousands of spatial locations in a tumor, but does not directly reveal genetic aberrations. We introduce CalicoST, an algorithm to simultaneously infer allele-specific copy number aberrations (CNAs) and a spatial model of tumor evolution from SRT of tumor slices. By modeling CNA-induced perturbations in both total and allele-specific gene expression, CalicoST identifies important types of CNAs – including copy-neutral loss of heterozygosity (CNLOH) and mirrored subclonal CNAs– that are invisible to total copy number analysis. On SRT data from nine patients from the Human Tumor Atlas Network (HTAN) with matched whole exome sequencing (WES) data, CalicoST achieves an average accuracy of 86%, approximately 21% higher than existing methods. On two patients with SRT data from multiple adjacent slices, CalicoST reconstructs a tumor phylogeography that describes the spread of cancerous clones in three-dimensional space. CalicoST analysis of multiple SRT slices from a cancerous prostate organ reveals five spatially coherent clones, with mirrored subclonal CNAs distinguishing clones on the two sides of the prostate, forming a bifurcating phylogeography in both genetic and physical space.

## 1 Introduction

Tumors evolve through acquisition of somatic mutations — including single nucleotide variations (SNVs), copy number aberrations (CNAs), and large-scale structural variations (SVs). Sequencing of somatic mutations in bulk tumors [1, 2] or disassociated single cells [3–5] has revealed the genetic heterogeneity within tumors and enabled the reconstruction of a tumor’s evolutionary history [6–8]. At the same time, tumors exhibit heterogeneity and undergo evolution within physical space, expanding and regressing based on interactions with other cells and the local microenvironment. Incorporating the spatial perspective into somatic evolution studies is a key challenge [9], but has been hampered by a lack of spatial data.

Recent technological advances in spatial sequencing provide a promising direction for studies of spatiotemporal tumor evolution. While high-quality spatial DNA sequencing would provide the ideal dataset for spatiotemporal evolution studies, such technologies remain in active development [10] and are not yet widely applied. However, spatially resolved transcriptomics (SRT) technologies that measure RNA simultaneously from thousands of spatial locations in a tissue has found extensive applications in analyzing the spatial organization of transcriptionally defined cell types within a tumor [11–14]. Even though somatic mutations occur in DNA and are not directly measured by SRT, large CNAs leave a signature in gene expression; namely, a deletion of a genomic region tends to result in underexpression of genes in the region, while an amplification tends to result in overexpression. Thus, identification of CNAs from transcriptomic data is a promising direction for analysis of somatic evolution in tumors.

Inferring CNAs from single-cell or spatially resolved transcriptomics is challenging as there are multiple explanations for an observed gene expression change, such as chromatin accessibility and transcription factor binding. It is typically difficult to determine whether an observed gene expression change is a result of CNAs or these other causes. Existing methods to infer CNA from gene expression data [15–17] assume that large CNAs alter the expression of multiple adjacent genes in a genomic region beyond expected by other regulatory effects. However, the variability of expression is so large between genes that these methods have limited accuracy in inferring CNAs and are not robust across tissues, patients, and cancer types. A few methods aim to address these challenges by combining single-cell RNA and DNA sequencing [18, 19], but few researchers perform both modalities of sequencing, limiting the usability of these methods. In addition, SRT, as well as single-cell RNA sequencing data (scRNA-seq) are sparse, generally having more than 75% zero counts across genes and cells/spots. Furthermore, SRT technologies (such as 10x Genomics Visium [20] and Slide-seqV2 [21]) pose an additional challenge beyond scRNA-seq: they measure a mixture of cells at each spatial spot, where normal cells can dilute the signals for CNAs.

Importantly, a CNA in cancer alters one of the two parental chromosomes, and thus the identification of *allele-specific CNAs* is essential for deriving a comprehensive description of CNAs in a tumor. For example, copy number neutral loss of heterozygosity (CNLOH) – an event where a region of one parental chromosome is deleted and the other parental chromosome is amplified so that the total copy number of the locus is unchanged – is common in cancer [22–24]. Similarly, mirrored-subclonal CNAs – where different cancer cells have independent gains or losses of different parental alleles – also occur in cancer [4, 25] but these events also result in identical total copy numbers across cancer cells. Thus, total copy number analysis does not reveal the complete CNA spectrum and may lead to incorrect identification of tumor clones and inaccurate tumor phylogenies. Previous methods have limited power to identify allele-specific CNAs in SRT data. Many existing transcriptomics methods cannot distinguish between alleles and only identify total changes in copy number [15–17]. A few methods for identification of allele-specific copy number aberrations from scRNA-seq data have been recently been developed [26–28], but these methods are challenged by the weak signal to distinguish the two parental alleles in scRNA-seq data.

We introduce a new method, CalicoST, that infers allele-specific CNAs in SRT data and uses these CNAs to reconstruct the phylogeographic evolution of a tumor. CalicoST identifies CNLOH and mirrored-subclonal CNAs that are invisible to total copy number analysis. CalicoST constructs a phylogeny of cancer clones that describes the accumulation of the inferred CNAs over time and a phylogeographic model that describes the spread of the tumor across physical space. We validate CalicoST using nine patients from the Human Tumor Atlas Network (HTAN) (WashU cohort) [29] with matched whole-exome sequencing (WES) data with high tumor purity. CalicoST achieves at least 86% accuracy in its inferred allele-specific copy numbers, 21% higher than previous methods. We reconstruct a three-dimensional phylogeography for a colorectal liver-metastasis and a breast cancer patients in HTAN with multiple adjacent slices. The phylogeography reveals the spatial direction of tumor growth, and particularly the expansion in the third dimension that could not be identified from a single slice. We also apply CalicoST to multiple SRT slices from a prostate cancer patient, identifying mirrored subclonal CNAs that suggest convergent evolution in the tumor. The reconstructed phylogeography partitions the cancer clones into the left and the right sides of the prostate, revealing the separation of the clones in both physical and genetic space. CalicoST enables the study of spatial tumor evolution, progression and metastasis, and will be helpful for further applications to cancer diagnosis and treatment.

## 2 Results

### 2.1 CalicoST algorithm

CalicoST infers allele-specific copy number aberrations (CNAs) and a phylogeographic model of tumor evolution from one or more SRT samples from a tumor (Fig. 1). CalicoST has the following key features.

**Figure 1:**
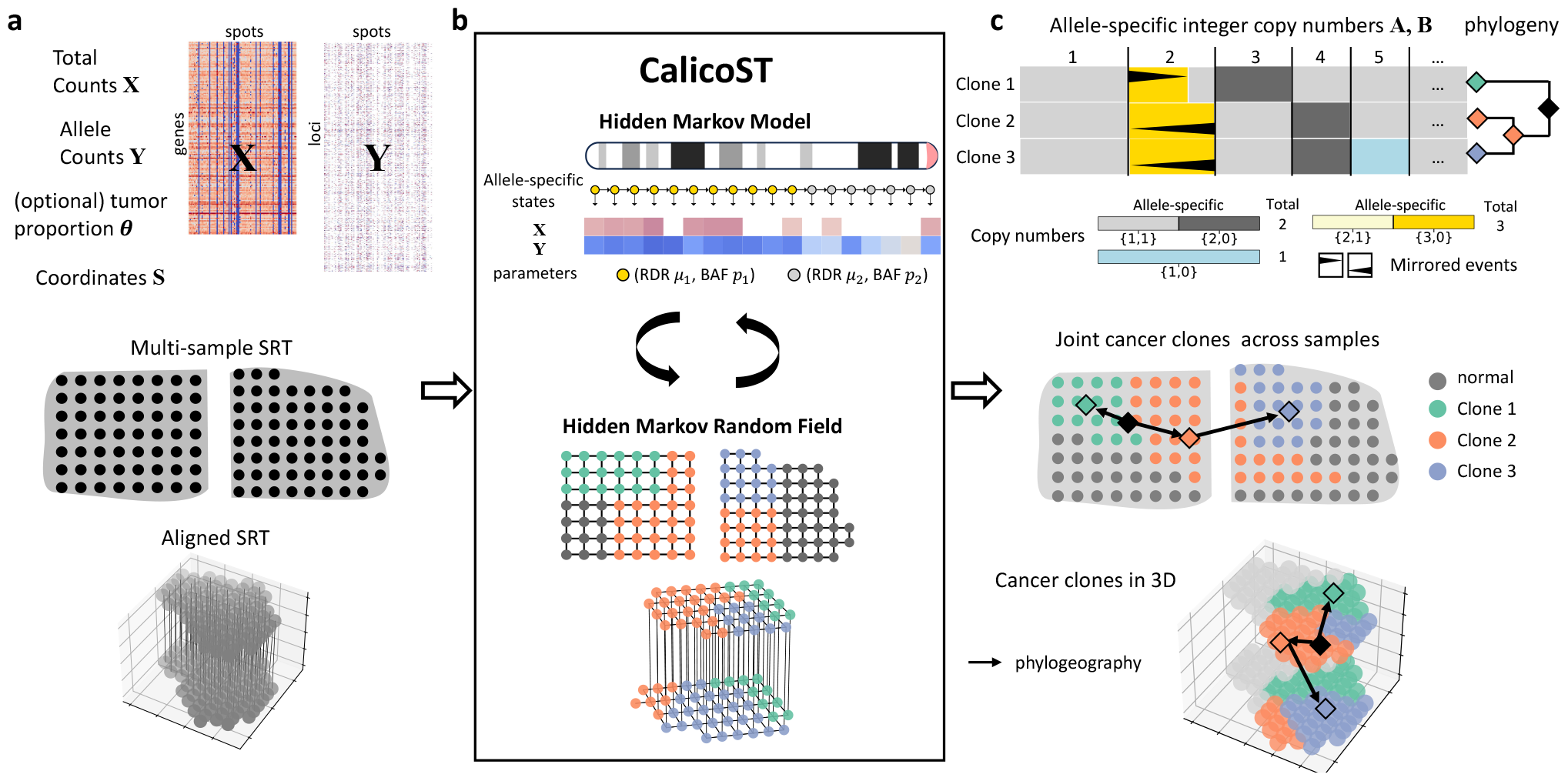
CalicoST infers allele-specific copy numbers and a phylogeography of a tumor from one or more SRT samples from the same patient. (a) Inputs to CalicoST are transcript counts **X**, allele counts **Y**, spatial coordinates **S**, and optionally the proportion of tumor counts per spot ***θ*** from one or more SRT slices or a 3D alignment of slices. (b) CalicoST jointly models transcript counts and allele counts as functions of allele-specific copy number states within each clone. CalicoST uses a Hidden Markov Model (HMM) to model correlations between copy number states from adjacent genomic regions and a Hidden Markov Random Field (HMRF) to model correlations between the cancer clones assigned to neighboring spatial locations. (c) CalicoST infers allele-specific integer copy numbers for one or more cancer clones, a phylogeny relating these clones, a clone label for each spot, and a phylogeographic model of the spatial expansion of cancer clones.

(1) Identifies allele-specific integer copy numbers for transcribed regions, revealing events such as copy neutral loss of heterozygosity (CNLOH) and mirrored subclonal CNAs that are invisible to total copy number analysis. (2) Assigns each spot a clone label indicating the clone it belongs to and the allele-specific copy number profiles it contain. (3) Infers a phylogeny relating the identified cancer clones as well as a phylogeography that combines genetic evolution and spatial dissemination of clones. (4) Handles normal cell admixture in SRT technologies that are not single-cell resolution (e.g. 10x Genomics Visium) to infer more accurate allele-specific copy numbers and cancer clones. (5) Simultaneously analyzes multiple regional or aligned SRT slices from the same tumor.

The inputs to CalicoST are a transcript count matrix **X** whose entries are the total number of reads from each transcript in each spot, and an allele count matrix **Y** whose entries are the number of reads from the non-reference allele of germline heterozygous SNPs (Fig. 1a). The matrix **X** is readily obtained from standard SRT analysis pipelines, while the matrix **Y** is calculated from a specialized pipeline that uses known locations of germline SNPs as well as reference-based phasing [30] to aggregate signal from multiple adjacent SNPs in the same haplotype (Section S3). This latter step is necessary because SRT data is generally sparse and the allele counts are even sparser: 98.8% SNP loci have zero total count within each individual spot and another 0.9% has only one total count.

In addition, some SRT technologies, (e.g. 10x Genomics Visium), may lack single-cell resolution, measuring multiple cells within each spatial spot. This admixture dilutes the signal for identification of CNAs and cancer clones. To ameliorate this issue, CalicoST optionally takes in a tumor proportion (***θ*** ∈ [0, 1]^*N*^ for *N* spots) as input. This proportion can be obtained using established methods for deconvolving cell type proportions in SRT data [31, 32].

The core of CalicoST is a generative probabilistic model of the observed variables **X, Y** as a function of the unobserved allele-specific copy numbers and clone labels 𝓁. Individual entries in **X** and **Y** provide poor estimates of the allele-specific copy number at the corresponding locus due to low sequence coverage and confounding by other sources of variation, such as variable gene expression. Thus, CalicoST aggregates signals from multiple adjacent loci in the genome and multiple adjacent spots. Specifically, we use a hidden Markov Model (HMM) to model correlations between the allele-specific copy number state of adjacent genomic loci and a hidden Markov Random Field (HMRF) to model the correlations between clone label 𝓁 in adjacent spots assuming that adjacent spots are likely to be genetically similar. We jointly infer the allele-specific copy number states and clone labels leveraging standard HMM and HMRF inference algorithms (Section 4.5,4.6).

Finally, we reconstruct a phylogeographic model to describe the ancestral relationships between the inferred clones as well as the spatial location of the ancestors of these clones (Fig. 1c). It is generally challenging to reconstruct a phylogeny from copy number profiles and requires complicated evolutionary models [33, 34]. Instead we leverage the fact that CalicoST infers loss of heterozyosity (LOH) events, which have the important property of being irreversible phylogenetic characters; i.e. once a parental haplotype is lost in a lineage, it cannot be regained. We construct a tumor phylogeny among cancer clones using the inferred LOH events and the star homoplasy model in Startle [35]. Then, we project the phylogeny in space and infer the spatial location of ancestors using a diffusion model (Section 4.7).

### 2.2 CalicoST infers accurate allele-specific integer copy numbers across HTAN samples

We evaluated CalicoST’s accuracy in inference of allele-specific copy numbers on 10x Genomics Visium Spatial Transcriptomics of twelve patients (twenty six slices) in HTAN (WashU cohort) [29] across three cancer types (Section 4.8). Whole exome sequencing (WES) data from adjacent bulk tumor sections was available for eleven patients. We determined the allele-specific integer copy numbers for nine bulk WES samples using HATCHet2 [36], while the remaining two samples have insufficient tumor purity to infer copy numbers. We used these copy numbers as the ground truth to benchmark the inferred CNAs from SRT data by CalicoST.

Across nine patients whose matched WES sample had sufficient tumor purity, the best-matching CalicoST cancer clone had 86% accuracy on average (min. 68% and max. 97%) (Fig. 2a), and an average of 95% precision and 90% recall in the prediction of genome segments with abnormal copy number (Section 4.9), respectively (Fig. S2a,b). The median length of CNA events detected by CalicoST was 80 Mb, often spanning entire chromosomes (Fig. 2b), which is of a lower resolution than CNA detected by HATCHet2 on WES samples. Nevertheless, CalicoST identified CNA events as small as 1 Mb for regions with high coverage. Notably, CalicoST infers allele-specific copy numbers and cancer clones from SRT data on two pancreatic cancer patients (HT270P1 and HT288P1) whose tumor purity in the bulk WES was insufficient for reliable identification of CNAs. We observed clear LOH regions from the observed B allele frequency (BAF) values of these two patients and the cancer clones are well distinguished by the read depth ratio (RDR) and BAF signals (Fig. S2c,d).

**Figure 2:**
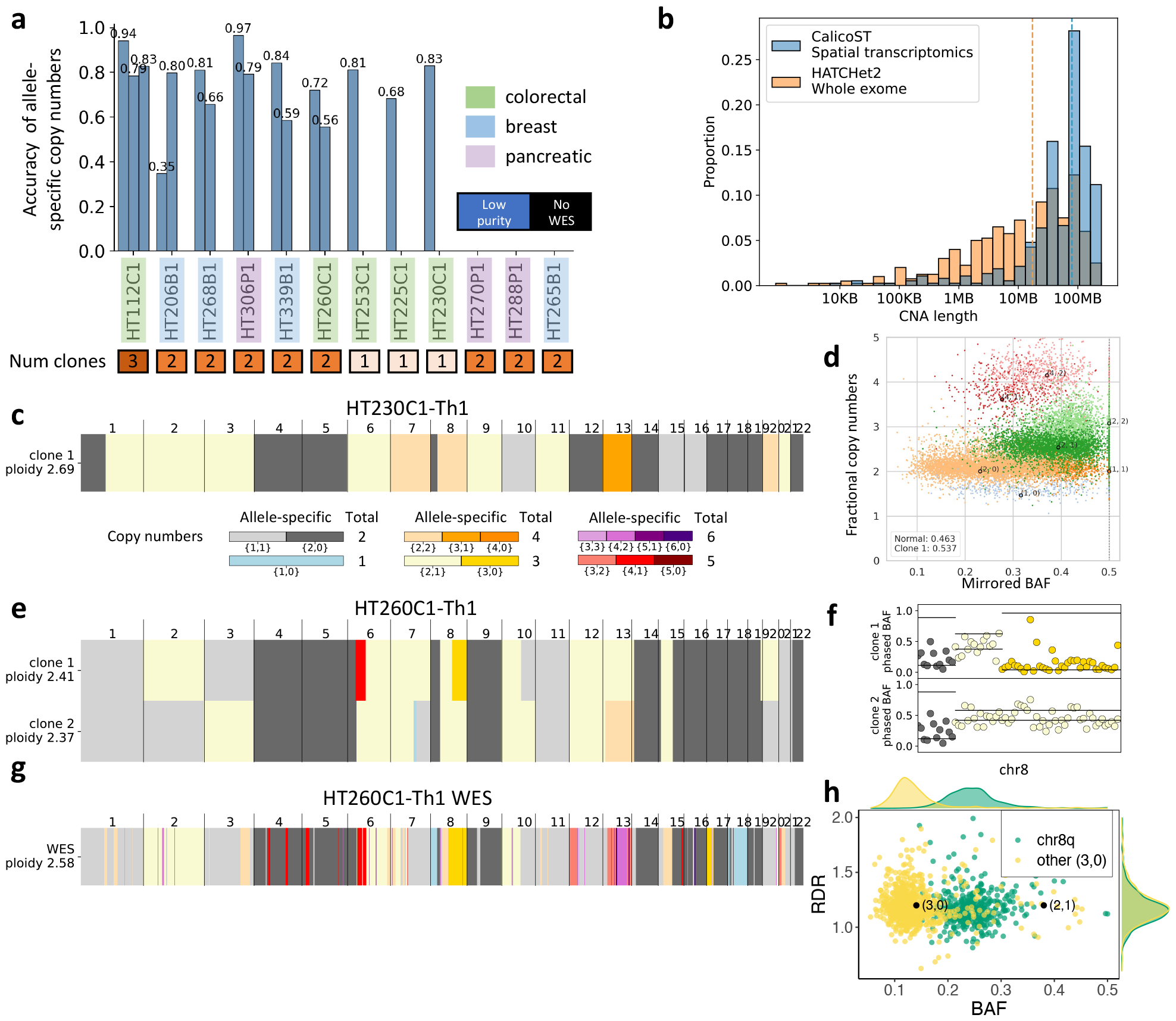
CalicoST infers accurate allele-specific copy numbers in HTAN samples. (a) Accuracy of allele-specific copy numbers across 12 HTAN patients (WashU cohort) inferred by CalicoST. Each bar represents an inferred cancer clone. (b) Length distribution of CNAs identified by CalicoST from SRT data and identified by HATCHet2 from WES for the nine patients with matched WES of sufficient tumor purity. The median length is 80Mb for CalicoST and 30Mb for HATCHet2 (vertical dashed lines). (c) Allele-specific integer copy numbers inferred by CalicoST from SRT data from CRLM patient HT230C1. Rows are cancer clones, and columns are genomic segments. Colors indicate allele-specific copy numbers. (d) The observed BAF (*x*-axis) and fractional copy numbers (*y*-axis) from the matched WES data of HT230C1-Th1. Each point is a genomic bin and colors indicate the allele-specific copy numbers inferred by HATCHet2 [36]. (e) Allele-specific integer copy numbers inferred by CalicoST from SRT data from CRLM patient HT260C1-Th1. (f) Observed RDR and BAF for chromosome 8 of HT260C1-Th1. Points are colored by the inferred allele-specific copy numbers. Horizontal black lines indicate the RDR and BAF of the corresponding copy number states estimated by HMM. (g) Allele-specific integer copy numbers inferred by HATCHet2 from WES from patient HT260C1-Th1. (h) The RDR and BAF values from WES data for bins from chromosome 8q region and bins from other genomic regions with {3,0} copy number state. Black points are expected RDR and BAF values for {3,0} and {2,1} states from HATCHet2 analysis.

CalicoST identifies large-scale changes in tumor ploidy that are often challenging to be inferred accurately, particularly for methods that infer only total copy numbers [36]. CalicoST identified near-triploid genomes in three colorectal liver-metastasis (CRLM) patients and a breast cancer patient (Fig. 2c,S3). For example, the estimated ploidy of CRLM patient HT230C1-Th1 is 2.7 with 40.5% and 12.6% of genomic bins having allele-specific copy number of {2, 1} and {2, 2} respectively (Fig. 2c). A triploid genome of this patient is also inferred in the matched WES sample (Fig. 2d). Previous studies have shown an association between triploidy and worse prognosis/poor survival [37]. Allelic information is key to the identification of neartriploid genomes from gene expression data; methods that only infer total changes in copy numbers miss many regions with copy number {2, 1} because the transcript counts in these regions many not differ substantially from copy number neutral regions, particularly because the gene expression signal is highly variable across the genome (Fig. S2e).

CalicoST also revealed tumor heterogeneity and clone-specific copy number alterations that were missed in bulk WES data. On CRLM patient HT260C1-Th1, CalicoST identified two cancer clones (Fig. 2e) with CNAs that were shared by both clones, such as CNLOH in chr17 and chr18, including well-known tumor suppressor genes *TP53* and *DCC* [38–41]. Other CNAs were unique to cancer clones. For instance, chr2 and chr3 had symmetric amplifications in the two cancer clones and chr8q has a loss of heterozygosity in clone 1 with allele-specific copy number {3, 0}, but has an allele-specific copy number of {2, 1} in clone 2. All three events are supported by the BAF signal in both clones (Fig. S4,Fig. 2f). The chr8q region was assigned to {3, 0} copy-number by HATCHet2 in the bulk WES (Fig. 2g). Although HATCHet2 detected one cancer clone in the bulk WES, its BAF and RDR measurements of chr8q highlight an unusual deviation from the expected BAF value of {3, 0} copy-number state (Fig. 2h), supporting CalicoST’s hypothesis that this region has undergone different CNAs in different cancer clones.

### 2.3 CalicoST identifies more accurate CNAs and spatially coherent clones than single-cell and spatial methods

We compared CalicoST with existing methods for identifying CNAs from single-cell RNA-sequencing (scRNA-seq) data [27, 42] and spatial transcriptomcs [17], evaluating both the accuracy of inferred CNAs and the spatial distribution of inferred cancer clones. Specifically, we compared CalicoST with (1) Numbat [16], an allele-specific CNA inference method for scRNA-seq data; (2) STARCH [17], a total copy number inference method for SRT data, and (3) inferCNV [42], a total copy number inference method for scRNA-seq data. Numbat and STARCH do not output integer copy numbers but rather copy number states (amplification, deletion, etc.), and thus their results are not directly comparable with CalicoST and HATCHet2. Thus, to perform a comparison, we projected the integer allele-specific copy numbers from CalicoST and HATCHet2 to copy number states (Section 4.9).

CalicoST had the highest accuracy on all but one sample, both when comparing allele-specific copy numbers to Numbat and total copy numbers to STARCH and inferCNV (Fig. 3a,S5). For allele-specific copy number, CalicoST was 25% more accurate than Numbat on average across the four samples. For total copy number, CalicoST had substantially higher accuracy in inferring CNAs than STARCH (59% higher on average) and InferCNV (90% higher on average). We compared the two allele-specific inference methods on all nine HTAN patients: CalicoST was 21% more accurate than Numbat on average and had better accuracy for eight of the nine patients (Fig. S5).

**Figure 3:**
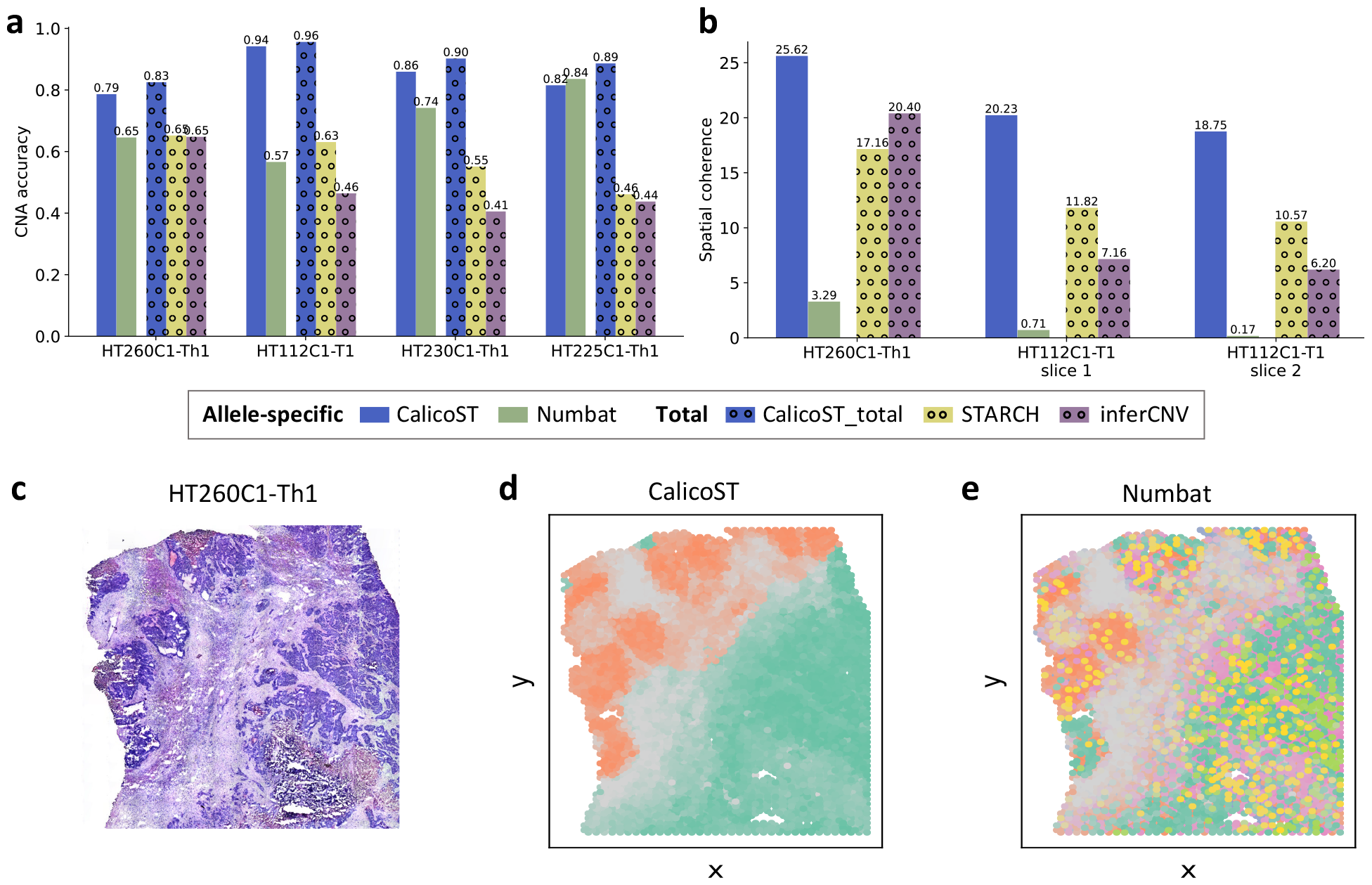
Comparing accuracy of CNAs and spatial coherence of inferred clones between Cali-coST and other CNA inference methods. (a) Accuracy and (b) spatial coherence comparison among CalicoST, Numbat, InferCNV, and STARCH on colorectal liver-metastasis (CRLM) patients. Solid bars indicate predictions of allele-specific copy number states and dotted bars indicate predictions of total copy number states. (c) H&E image of a CRLM sample HT260C1-Th1. (d) Cancer clones inferred by CalicoST. *x*- and *y*-axes are spatial coordinates, and grayscale represents the proportion of normal cells within each spot, as inferred by RCTD. Other colors indicates cancer clones. (e) Cancer clones inferred by Numbat using the same color scheme as (d).

The spatial distribution of cancer clones inferred by CalicoST was substantially more coherent than the other three methods on the two CRLM patients where all methods identify multiple cancer clones (Fig. 3b, Fig. S5). For example, on patient HT260C1-Th1 (Fig. 3c), CalicoST identified two spatially coherent clones that partition the tissue into the top left and bottom left regions, partitioned by normal spots indicated by the gray color (Fig. 3d). In contrast, Numbat identified cancer clones with lower spatial coherence with some cancer clones (yellow and pink) spread almost uniformly through the slice and on both tumor regions separated by normal spots (Fig. 3e).

### 2.4 CalicoST reconstructs tumor evolution in three-dimensional space

We applied CalicoST to infer phylogeographic trees in space (phylogeography in short) for two HTAN patients where 10x Genomics Visium Spatial Transcriptomics data was obtained from multiple adjacent slices of the tumor: CRLM patient HT112C1-T1 with two slices separated by 60 µm and breast cancer patient HT268B1-Th1 with five slices, with a distance of 100 µm between four of the slices, and an unknown distance between the first two slices. We aligned adjacent sections and derived a multi-slice alignment using PASTE2 [43], which was input into CalicoST.

For the CRLM patient, CalicoST identified three spatially coherent clones in the 3D tumor tissue and infers a phylogenetic tree from the CNAs in these clones (Fig. 4a). This phylogeographic tree shows the expansion of the tumor, branching from the ancestral clone 1 (green) to two clones on either side (orange and blue). The three clones have distinct allele-specific copy number profiles (Fig. 4b). Specifically, the orange clone 2 has a unique LOH on chr21, and the blue clone 3 has a unique LOH on chr 11p. Both events are supported by a strong allelic imbalance in the BAF (Fig. S6a). We observe a high consistency in clone composition and localization between the two slices, which is not surprising as the distance between the two slices (60 µm) is small and almost the same as the diameter of a spot within one slice (55 µm).

**Figure 4:**
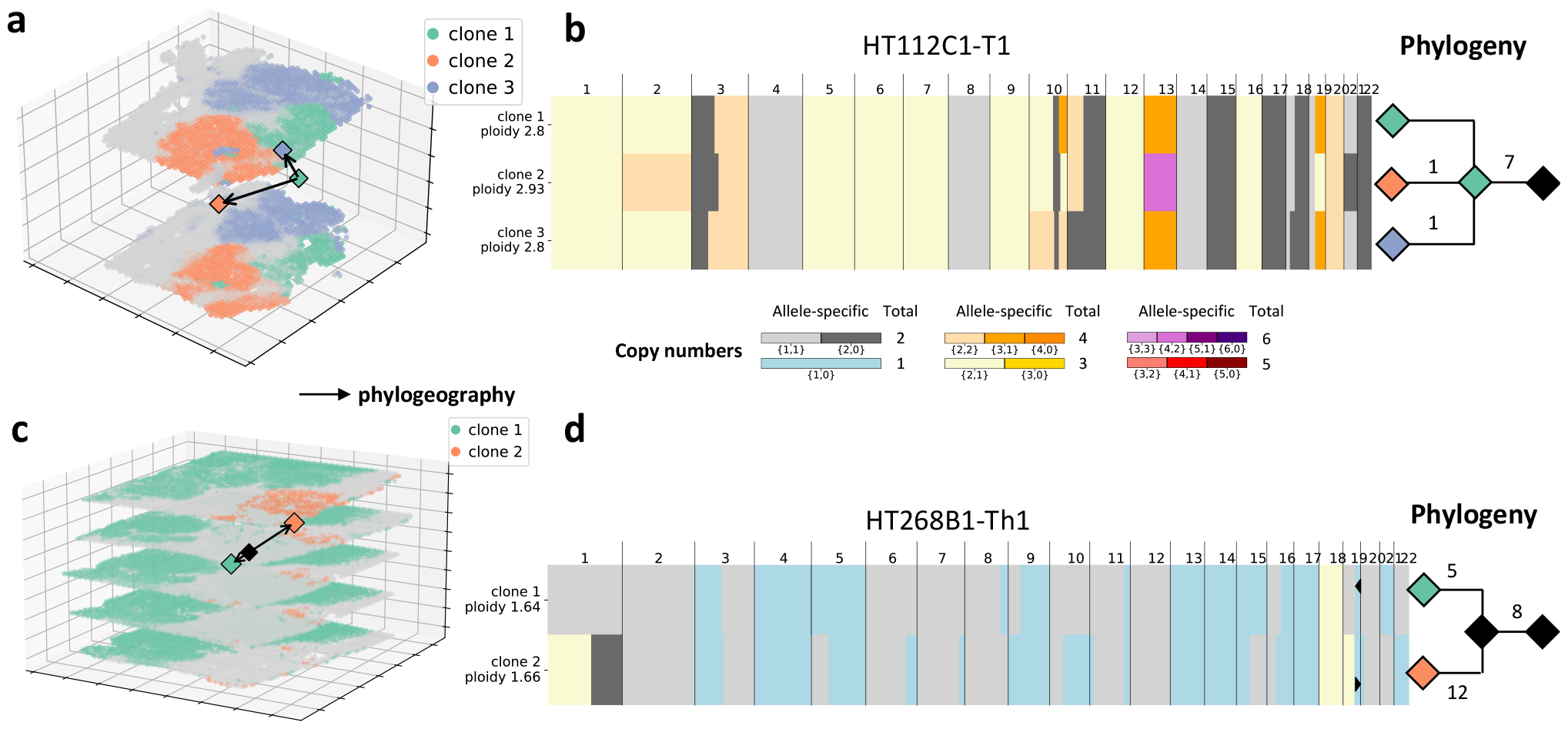
Tumor evolution in 3D inferred by CalicoST from patient HT112C1-T1 and HT268B1-Th1. (a) Spatial distribution of three cancer clones identified by CalicoST in two adjacent slices from CRLM patient HT112C1-T1. Grayscale indicates the inferred proportion of normal cells within each spot. Diamonds are the spatial centroid of each clone or inferred ancestor locations and arrows indicate the inferred directions of tumor development. Distance between two slices in the *z* coordinate is enlarged for clearer visualization. (b) Allele-specific copy number profiles for the three cancer clones and the corresponding phylogeny (right) with branches in the phylogeny labeled by the number of unique large LOH events that occur on the branch. (c) Spatial distribution and phylogeographic tree of two cancer clones in five adjacent slices from breast cancer patient HT268C1-Th1. Color scheme is the same as (a). (d) Inferred allele-specific copy numbers and reconstructed tumor phylogeny.

CalicoST identifies two cancer clones in a breast cancer patient HT268B1-Th1 across five slices that are aligned in 3D space and reconstructs a phylogeography between the two clones (Fig. 4c). The phylogeography indicates the ancestor (black diamond) is located between the two clones, which expanded leftward and downward along the *z* axis to clone 1 (green) and rightward and upward to clone 2 (orange). The spatial evolution of this patient contains a strong component in the *z*-axis direction, which can only be revealed due to the multiple slices of SRT data. The two clones have copy number aberrations that are shared between both clones and unique to each clone including a mirrored deletion on chr19 (Fig. 4d), which is supported by the RDR and BAF values in this genomic region (Fig. S6b). Notably, clone 1 has fewer unique LOH events than clone 2, suggesting that clone 1 is genetically closer to the common ancestor than clone 2, which is reflected in the inferred location of the ancestor in the phylogeography (Fig. 4c).

### 2.5 Mirrored copy number aberrations in multiple regions of a cancerous prostate organ

We applied CalicoST to infer allele-specific CNAs and a phylogeography jointly from five slices from a single cross-section of a cancerous prostate [44] (Section 4.8). CalicoST identifies five cancer clones across the SRT slices with some clones shared across multiple slices. (Fig. 5a). The spatial distribution of the inferred cancer clones are visually consistent with the pathologist-annotated tumor regions shown in [44], even though CalicoST was not given information about the locations of normal spots or estimated tumor purity in each spot. The five clones have distinct copy number profiles (Fig. 5b), which are supported by the BAF in each clone (Fig. S7). Notably, clone 5 (blue) is shared across all three slices on the right side of the prostate organ, forming a contiguous spatial region, even though CalicoST was not given information about the relative locations of slices in the prostate. This demonstrates the advantage of CalicoST’s joint inference across multiple slices.

**Figure 5:**
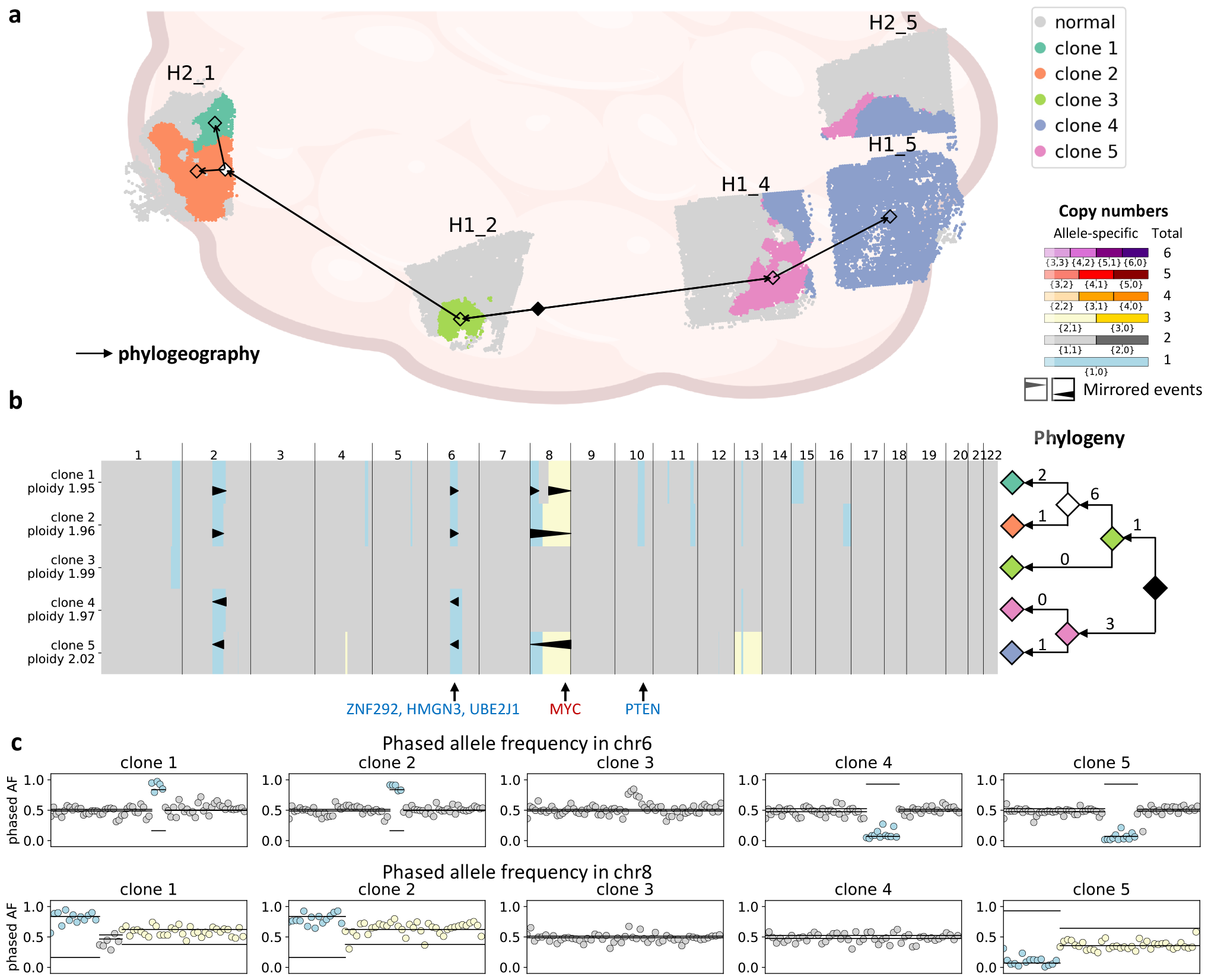
CalicoST infers a phylogeography and mirrored CNA events in a cancerous prostate. (a) Spatial distribution of cancer clones inferred jointly by CalicoST across five slices from a cancerous prostate. Positioning of five slices is according to [44]. Colors indicate inferred clones, including the normal clone in grey. Arrows represent the phylogeography of tumor evolution. (b) Allele-specific copy number profiles for the five cancer clones and the corresponding phylogeny (right) with branches in the phylogeny labeled by the number of unique large LOH events that occur on the branch. Colors indicate allele-specific copy numbers. The orientation and position of triangles indicate mirrored CNA events. (c) BAF of each clone in (top) chr6 and (bottom) chr8. Colors indicate allele-specific copy numbers using the same color scheme as panel (b).

The clones on the two halves of the prostate cross-section (left and right) are distinguished by multiple aberrations that are unique to each half. Most prominent among these are four mirrored CNA events, on chromosomes 2, 6, and 8 where clones 1 and 2 have different amplified/deleted alleles than clones 4 and 5 (Fig. 5c). Specifically, chr8p, one of the most frequently deleted regions in prostate cancer [45], has a mirrored deletion in the two halves of the prostate, and chr8q has a mirrored amplification containing the *MYC* gene, a well-known oncogene in aggressive prostate cancer. Also, chr6 has a mirrored deletion of the region 73-99Mb, which is also a commonly deleted region in prostate cancer [46], harboring three reported tumor suppressor genes (*ZNF292, HMGN3*, and *UBE2J1*) [47]. The occurrences of three independent deletions and an independent amplification in different clones indicate of the high frequency of corresponding events in prostate cancer and are potential signal of convergent evolution [25]. In contrast, the original published analysis of this data [44] used InferCNV, and concluded that the deletion in chr6q is a truncal event based on changes in total copy numbers, missing the differential loss of the two alleles in the two halves of the prostate.

The inferred phylogeography splits the cancer clones into two main lineages, which coincide with the left and right spatial partition of the prostate cross-section (Fig. 5a). On the left half, clone 3 contains only one CNA event, a deletion in chr1, and is the closest to a normal state; clones 1 and 2 share the deletion in chr1 and many other deletions in chromosomes 2, 4, 6, 8, 10, and 11. On the right half, clone 4 shares multiple CNAs with clone 5, but does not have any unique CNA and thus clone 4 is marked as an ancestor of clone 5, which is consistent with its spatial location closest to the root. Interestingly, the absence of truncal CNA events and the clear bifurcation in both genetic and physical space suggests that the tumor on the left and right halves diverged in a very early stage and had relatively independent evolution.

## 3 Discussion

We introduced CalicoST, an algorithm that infers allele-specific copy numbers and reconstructs a phylogeography relating cancer clones in time and space using SRT data. We applied CalicoST to SRT data from twelve HTAN patients across three cancer types (WashU cohort) and multiple slices from a cancerous prostate. CalicoST showed high concordance with copy number aberrations (CNAs) identified in bulk whole-exome sequencing (WES) from nine patients with sufficient tumor purity, but also revealed multiple cancer clones in many samples as well as cancer clones in low purity samples that were not identified in bulk WES. CalicoST is more accurate and yields more spatially coherent clonal organization compared to existing methods that identify CNAs from single-cell or spatial transcriptomics data.

CalicoST uses the inferred CNAs to construct a phylogeny relating the cancer clones, and the spatial locations of the cancer clones to construct a phylogeography, which combines both the genetic and spatial evolution of a tumor in a unified model. This reconstruction is enabled by allele-specific copy numbers, as CalicoST uses loss of an allele/haplotype as irreversible phylogenetic characters, circumventing some of the difficulties in deriving phylogenies from copy number aberrations [33]. Applied to colorectal liver-metastasis samples from HTAN with multiple consecutive slices, we construct 3D models of spatial tumor evolution which describe both the genetic aberrations and spatial directions of tumor growth. On a spatial transcriptomics dataset containing multiple sections from cancerous prostate [44], CalicoST identifies mirrored subclonal copy number aberrations that are missed in the analysis of total copy number; moreover, CalicoST infers a phylogeography that bifurcates the left and right halves of the prostate in both genetic and physical space, pointing toward an early divergence between tumor cells on different halves of the prostate.

CalicoST has some limitations, some of which are directions for future improvement. First, the length of the copy number aberrations that can be reliably detected is limited by the sequencing coverage as well as the inherent difficulties in detecting DNA aberrations from gene expression data. On the 10x Genomics Visium data analyzed in this study, we detected aberrations as small as 1 Mb, with a median aberration size of 80 Mb. This resolution depends on the gene density within a genomic region as well as the number of spots that contain the aberration. However, it will be nearly impossible to detect aberrations in single genes, since these are indistinguishable from differential expression. Second, CalicoST’s use of allele/haplotype deletion as phylogenetic markers helps infer accurate phylogenies, but requires that a tumor sample have enough of these events. Some tumors may have insufficient losses to yield robust phylogenies, particularly among tumors containing many cancer clones. Leveraging other CNAs in phylogeny reconstruction may address this issue but requires further investigation of the trade-off between the increased number of events and potential inaccuracies in phylogeny inference. Third, CalicoST struggles with inferring the exact integer copy numbers for amplifications with a high total copy number because of the high variance in gene expression. For example, CalicoST infers chr13 of HTAN patient HT260C1-Th1 to have three and four total copies across the inferred cancer clones but the total copy numbers inferred from WES data by HATCHet2 is five copies. Inference of CNAs jointly from SRT and DNA data may help with this issue, when both measurements are available. Fourth, further improvements can be made in the model selection criteria that CalicoST uses to select the number of clones (Section S7) and the parameter in the HMRF that governs the spatial coherence of the inferred clones (Section 4.6). Particularly, for tumor samples containing cancerous cells with little spatial organization, a strong spatial coherence prior may lead to inaccurate inference of CNAs and cancer clones.

The use of spatially resolved transcriptomics in cancer analysis is growing rapidly. CalicoST can help bring valuable insights into copy number drivers of cancer, spatial tumor heterogeneity, and spatial evolution, serving as a foundation for additional biological analyses integrating genetic evolution, epigenetic (gene expression) changes, and spatial organization.

## 4 Methods

### 4.1 CalicoST workflow

CalicoST has the following main steps. In a preliminary step, CalicoST extracts allele and total counts at germline heterozygous SNP loci to distinguish between alleles (Fig 6 step 0, Section S1). The first step is to aggregate counts along genome to reduce the sparsity (Fig 6 step 1, Section S3). Because each CNA event occurs on a parental allele, we infer a grouping of SNPs by parental alleles (also known as phasing) to avoid mixing the counts. The second step is to infer normal spots (Fig 6 step 2, Section S4). The inferred normal spots provide baseline gene expression in normal cells; higher-than-baseline expression are potentially due to copy number gains and lower-than-baseline expression indicate copy number losses. Additionally, we use normal spots to remove genomic bins that potentially have allele-specific gene expression irrelevant to CNAs (Section S5). In the third step, CalicoST relaxes the constraint that allele-specific copy numbers are integers, and clusters genomic bins into *copy number states* and infers cancer clones simultaneously (Fig 6 step 3). CalicoST explicitly models the correlation among genomic bins along the genome using a Hidden Markov Model and among the cancer clones in space using a Hidden Markov Random Field in this step. Particularly, CalicoST estimates a latent value for the read depth ratio (RDR) in HMM underlying the genomic bins corresponding to each copy number state to indicate the relative copy numbers compared to diploid, and a latent value for the BAF for each copy number state to indicate the imbalance of copy numbers between the two alleles. Next, CalicoST finds allele-specific integer copy numbers for each copy number state that best explain the inferred latent RDR and BAF (Fig 6 step 4, Section S6). Finally, Cali-coST reconstructs a tumor phylogeography by inferring a phylogeny using the inferred LOH events and projecting to space.

**Figure 6:**
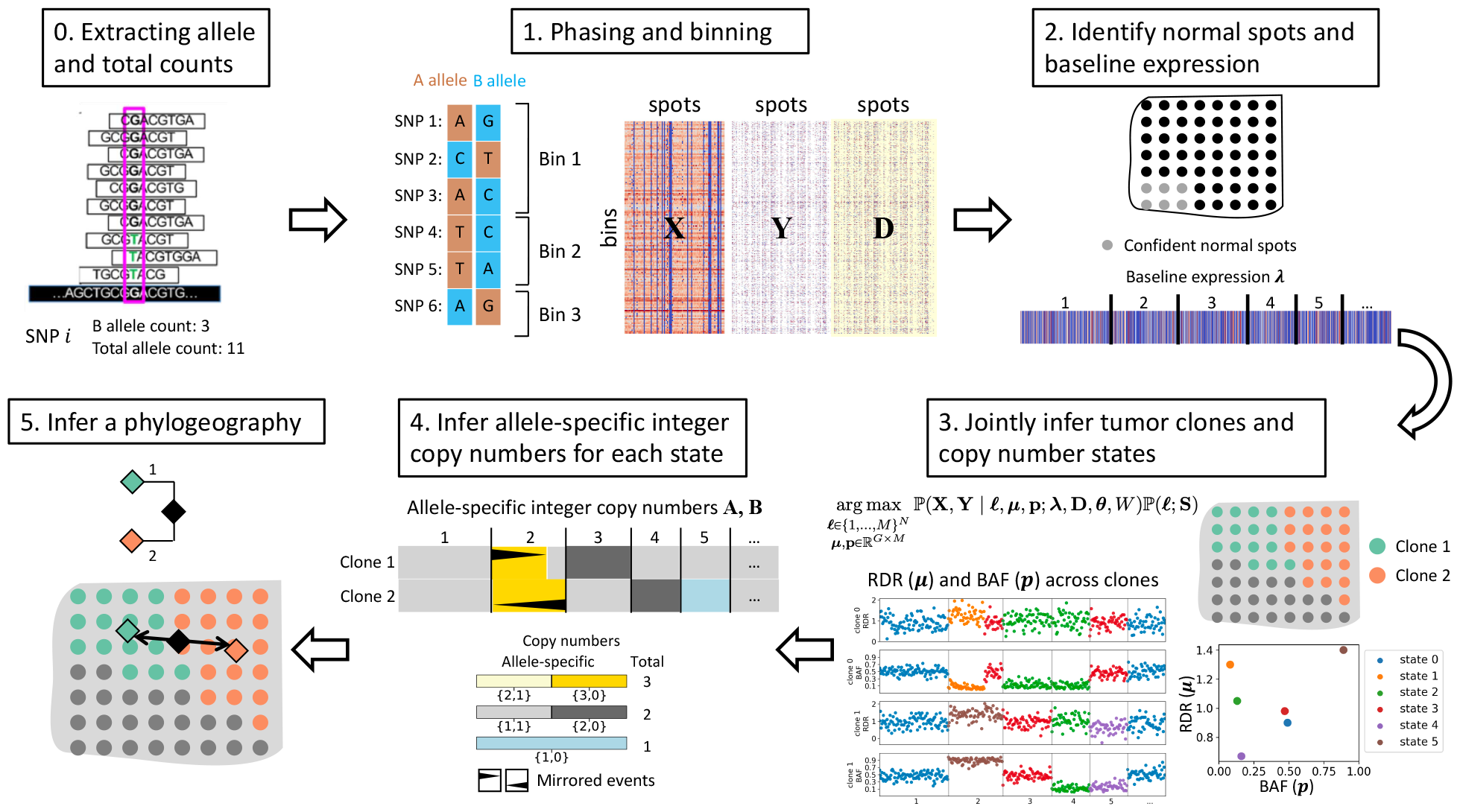
Workflow of CalicoST. CalicoST extracts the B allele and total counts of heterozygous SNP loci from the BAM file to distinguish between two alleles. CalicoST phases the SNPs and aggregates transcript counts and allele counts of each haplotype along the genome. Next, CalicoST identifies normal spots, jointly infers tumor clones and estimates a latent RDR and BAF value for each copy number state across all involved genomic bins in each clone. CalicoST infers integer allele-specific copy numbers using the latent RDR and BAF values. Finally, CalicoST reconstructs a phylogeographic model of tumor evolution by inferring a tumor phylogeny using LOH events and inferring spatial locations of ancestral clones.

In the sections below we provide further details for steps 3 and 5 of CalicoST. Section 4.2 describes the objective function of inferring allele-specific CNAs. Section 4.3 explains the underlying probabilistic model of the observed counts. Section 4.4, 4.5, and 4.6 describe the solution of the allele-specific CNA inference objective function. Section 4.7 explains the phylogeography reconstruction. The remaining subsections describe the details of analyses on HTAN (WashU cohort) and the prostate cancer samples.

### 4.2 Copy number aberrations (CNAs) and clone inference problem

Given the aggregated transcript counts **X** = [*x*_*g,n*_], phased B allele counts **Y** = [*y*_*g,n*_], and total allele counts **D** = [*d*_*g,n*_] across *n* = 1, …, *N* spots and *g* = 1, …, *G* genomic segments, CalicoST finds a clone label 𝓁 ∈ {1, …, *M*}^*N*^ to indicate one of the *M* clones each spot belongs to, and two allele-specific copy number matrices for each clone for each segment, **A** = [*a*_*g,m*_] of A allele copies, and **B** = [*b*_*g,m*_] of B allele copies.

CalicoST formulates a maximum likelihood problem to infer 𝓁, **A** and **B**. CalicoST also uses the following quantities in the problem: the normalized transcript counts in normal cells 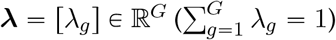, and the spatial coordinates **S** ∈ ℝ^*N*×2^. CalicoST optionally takes in the alignment *W* across slices when jointly identifying clones and CNAs across multiple SRT slices, and the *tumor count proportion θ* = [*θ*_*n*_] ∈ [0, 1]^*N*^ for each spot *n*. The overall likelihood objective of CalicoST is:

#### CNA and clone inference problem

*Given SRT data* (**X, Y, D, S**), *optionally* (*θ, W*), *and a given the number M of clones, find clone labels* 𝓁 *and integer allele-specific copy numbers* **A** *and* **B** *that maximize the log-likelihood of the data:*

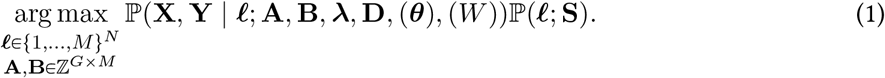

Solving this problem with integer-valued **A** and **B** is challenging. Notably, the probabilistic model of **X** involves fractional values derived from all values in **A** and **B**, as detailed in the following section. Previous work on copy number inference [4, 28, 36, 48] usually transform the integer copy numbers to a discrete set of real-valued latent parameters: *read depth ratio (RDR)* ***µ*** and *B allele frequency (BAF)* **p**. We use the same parameter transformation and split the problem into step 3 to infer clone labels and the latent RDR and BAF parameters and step 4 to infer **A, B** from the estimated RDR and BAF parameters.

Allele-specific copy numbers can only take values from a finite set of size *K*, which we call the *copy number states*. Accordingly, the latent RDR and BAF also have *K* unique values. We introduce a categorical variable **Z** = [*z*_*g,m*_] ∈ {1, …, *K*}^*G*×*M*^ to indicate which of the *K* copy number states each genome segment in each clone takes. We infer *K* RDR parameters *µ* = [*µ*_*k*_] ∈ ℝ^*K*^, *K* BAF parameters **p** = [*p*_*k*_] ∈ ℝ^*K*^, state indicator **Z**, and clone labels 𝓁 by:

#### Copy number state and clone inference problem

*Given SRT data* (**S, X, Y, D**), *optionally* (*θ, W*), *and a given the number M of clones, find clone labels* 𝓁, *copy number states* **Z**, *latent RDR* ***µ*** *and BAF* **p** *that maximize the log-likelihood of the data:*

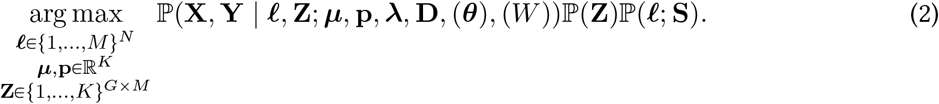

We model the correlation of copy number states along the genome by specifying ℙ (**Z**) as a Markov chain, and the correlation of cancer clones in space by specifying ℙ(𝓁) as a Markov Random Field. Overall, the likelihood (2) combines a Hidden Markov Model of copy number states with a Hiddem Markov Random Field of cancer clones. In the notation of probabilistic models, we separate the conditional random variables from the parameters and constants by a semicolon. So 𝓁 and **Z** are random variables in the above equation, and ***µ*, p, D**, *θ* and *W* are parameters or constants. We also put the optional input data in parentheses.

### 4.3 Copy number probabilistic model

Copy number aberrations affect **X** and **Y** in the following ways: increasing total copy number leads to increased gene expression and hence higher values in the corresponding entries in **X**; increase or decrease of copy number of one allele leads to imbalanced read counts between the two alleles and hence the ratio between **Y** and **D** is biased away from 0.5 at corresponding entries. Considering that spatial spots contain a mixture of tumor and normal cells, the degree of increased gene expression or imbalanced alleles depends on the proportion of reads coming from tumor (or normal). We derive the probabilistic model when assuming each spot contains a homogeneous tumor clone in the following of this section, and extend to the case of a tumor-normal mixture in Section S2.

Let 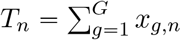 be the total transcript counts across all genomic bins for the *n*^*th*^ spot. Suppose the cells in this spot are all from clone *m*. We assume the copy numbers at each bin *g* scale the baseline proportion of transcript counts *λ*_*g*_ by 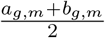. We model *x*_*g,n*_ by a Negative Binomial distribution parameterized by *T*_*n*_, ***λ*, A, B** and an additional over-dispersion parameter *ϕ*:

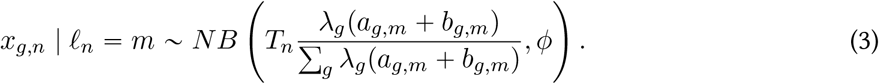

The Negative Binomial distribution can be viewed as an approximation for the Dirichlet Multinomial distribution *DirMult*(*T*_*n*_, *α*), where 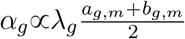. The Multinomial probability parameters are constrained to be a simplex, and are more difficult to optimize than a Negative Binomial distribution.

We model **Y** using a Beta-binomial distribution given the ratio between B allele copy number and total copy numbers at each genomic bin and the total SNP-covering reads **D**. The Beta distribution prior in the Beta-binomial distribution allows large variance than a binomial distribution, thus taking into account potential sequencing biases and other unknown factors related to allele imbalance.

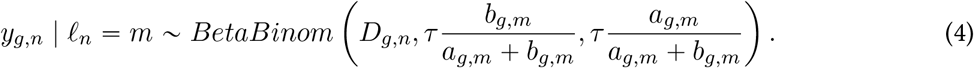

The mean of the Negative Binomial distribution has a fractional form of **A, B**, making a direct optimization of integer **A** and **B** challenging. We transform **A** and **B** into latent RDR ***µ*** and BAF **p** parameters with *K* unique values across copy number states. Suppose the segment *g* in clone *m* takes the *k*^*th*^ copy number state, the corresponding latent RDR and BAF is:

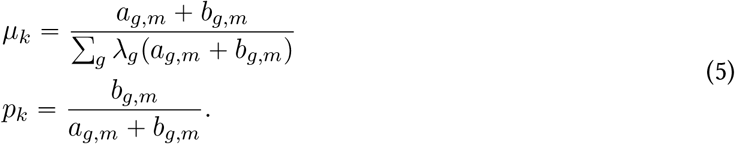

Note that the denominator of *µ*_*k*_ is a weighted average of total copy numbers along the genome in each clone, and technically takes different values in different clones. But under the assumption that different clones share many common CNA events, specifically when they are close in lineage, we assume the denominators are similar across clones and the *K* unique RDR values ***µ*** are shared across clones. This transformation (5) is the basis for inferring integer copy numbers (Section S6). Also note that we express the probabilistic model for individual spots, but it is generalizable to a pseudobulk of multiple spots.

#### Visualization the data

We define *observed RDR* as 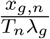 and *observed BAF* as 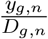 for data visualization. Additionally, if the latent RDR and BAF are found to be close to 1 and 0.5, respectively, along the genome in a clone, we drop this clone for visualization because it does not contain CNAs detectable to CalicoST and likely contains mainly normal cells.

### 4.4 Block coordinate ascent optimization of joint likelihood

The clone labels 𝓁 and the copy number states and parameters **Z, *µ*, p** are interleaved in the probabilistic models (3)(4). To make the optimization tractable, we use a block coordinate ascent method to solve for 𝓁 and for ***µ*, p, Z** iteratively. Given 𝓁, we solve for ***µ*, p, Z** under the Hidden Markov Model in Section 4.5; given ***µ*, p, Z**, we solve for 𝓁 under the Hidden Markov Random Field in Section 4.6.

### 4.5 Hidden Markov Model (HMM) to infer copy number states

Given clone labels 𝓁, we optimize the following objective for **Z, *µ***, and **p**:

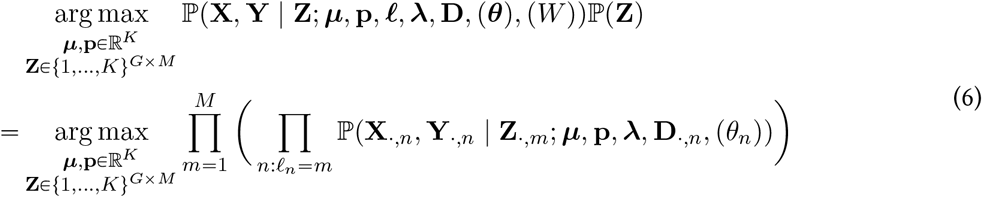

With an abuse of notation, we use 𝓁 to denote the values that the random variable of clone label takes, rather than the random variable itself.

Given that CNAs affect large contiguous regions in the genome, adjacent genomic bins tend to have the same copy number state. We model the copy number states **Z**_·,*m*_ using a Markov model for each clone *m* with equal values for the start probability and inter-state transition probabilities:

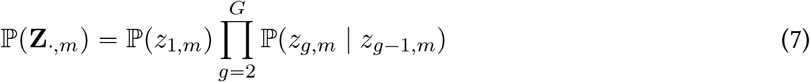

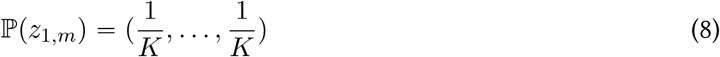

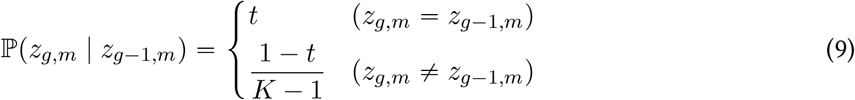

where the parameter *t* is a user-defined parameter of self-transition probablity. The objective (6) is a hidden Markov Model (HMM) under this prior distribution of ℙ (**Z**_·,*m*_). While the transition probability *t* can be inferred during HMM inference, **X** and **Y** tend to have large variances in SRT data; thus, the estimated *t* tends to favor a high probability of inter-state transition and disagrees with CNA event sizes and frequencies in reality. We use *t* = 1 − 10^−5^ by default. We use the Baum-Welch algorithm to estimate RDR *µ* and BAF **p** parameters.

While the MLE estimate of **Z** in (6) can be solved by Viterbi algorithm, we instead compute the full posterior distribution of *z*_*g,m*_ given by the forward-backward algorithm, which marginalizes *z*_*g,m*_ over all possible copy number states of other segments.

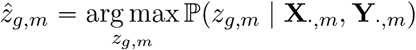

In practice, the counts of individual spots are still sparse and the HMM likelihood optimization may fall into local maxima. We showed that the likelihood of aggregated counts across spots within each clone is only different from that of individual spots by a constant, if dropping the over-dispersion parameters in the probabilistic model (Section S9). But the aggregated counts are much less sparse and suffer less from the local maxima, therefore we optimize the following likelihood function.

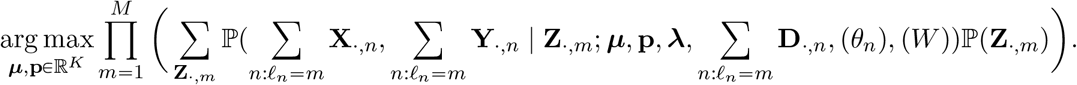

### 4.6 Leveraging spatial coherence for inferring clone labels by Hidden Markov Random Field

Given an estimated RDR *µ* and BAF **p** and the most probable values of **Z**, we optimize the following objective over clone labels 𝓁

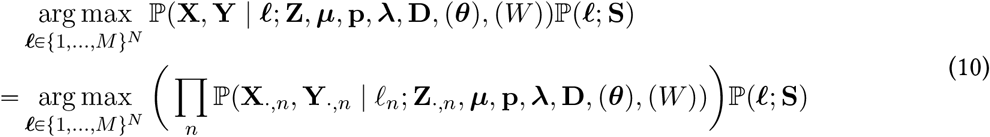

We assume clones are spatially coherent and impose a Potts model [49] as the prior distribution for ℙ(𝓁; **S**). Let 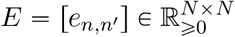 be the weighted spatial adjacency matrix, which combines intra-slice adjacency and inter-slice alignment *W*. Let *α*_*m*_ be the proportion of spots in clone *m* and let *β* be the strength of spatial coherence. Then

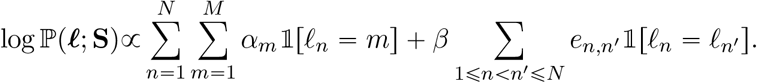

Note that the weighted adjacency matrix *E* contains both within-slice spatial adjacency and the alignment *W* across slices if it is available. The objective (10) is a hidden Markov Random Field (HMRF) and we use iterated conditional modes [50] for optimizing 𝓁.

Notice that we obtain the full posterior probability of ℙ(*z*_*g,m*_ | **X**_·,*m*_, **Y**_·,*m*_) via forward-backward algorithm, and accordingly we give the option in CalicoST to leverage the full posterior probability. Denote the full posterior probability ℚ(**Z**_·,*m*_) =Π_*g*_ ℙ(*z*_*g,m*_ | **X**_·,*m*_, **Y**_·,*m*_), CalicoST can alternatively solve the following objective function that uses ℚ(**Z**_·,*m*_):

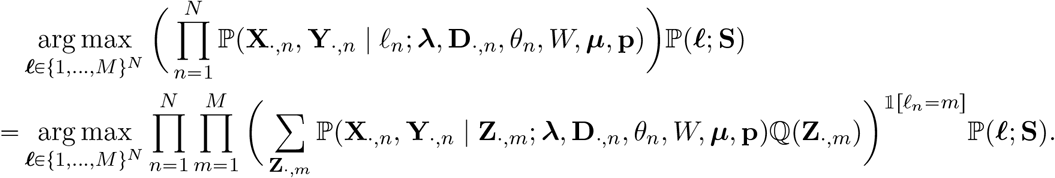

### 4.7 Reconstructing tumor phylogeography

We use a two-step approach to reconstruct a tumor phylogeography: first inferring a tumor phylogeny using the inferred CNAs and then projecting the tumor phylogeny in space and inferring ancestor spatial locations.

We apply Startle [35] to reconstruct a tumor phylogenetic tree among CalicoST-inferred clones using the inferred LOH events. Using LOH events in phylogeny reconstruction brings the following advantages: firstly, LOH events can be more accurately identified by the imbalanced BAF signals than other CNAs; secondly, LOH is irreversible when traversing the phylogenetic tree from the root to each leaf, and thus compatible with the state-of-the-art phylogeny reconstruction methods such as Startle. Startle infers a phylogeny to describe how the llabels” of a list of genomic lsites” evolve. The list of lsites” is the refinement of genome partitions based on CNAs in our application, and the llabels” are one of the three states: no LOH, LOH of A allele, and LOH of B allele. Startle finds a phylogenetic tree with minimum number of LOH events along the edges.

To infer a phylogeography, we project the leaf nodes (which correspond to inferred clones) to the center of involved spots in space (denoted by *s*_*v*_ for node *v*), and infer the spatial location of ancestor nodes using a Gaussian diffusion model. Specifically, we assume the spatial distance between a node *v* and its parent *p*(*v*) in the phylogenetic tree follows a Gaussian distribution with a variance proportional to the number of mutations *wv,p*(*v*) on the edge.

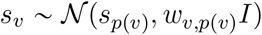

We estimate the ancestor locations in the phylogenetic tree by maximizing the joint probability of spatial locations of all nodes, {*s*_*v*_}_*v*_, under the above Gaussian distribution:

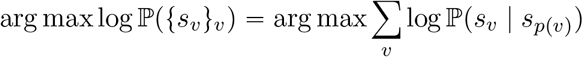

### 4.8 Running CalicoST on HTAN and prostate cancer samples

For each HTAN patient, we used CalicoST to infer CNAs and tumor clones jointly across all SRT slices. If a patient has multiple slices, we jointly identified germline heterozygous SNPs across all slices before running CalicoST to increase the SNP calling sensitivity. We ran CalicoST with the tumor count proportions *θ* that are derived from deconvolving SRT spots using matched and cell-type-annotated snRNA-seq data using RCTD [31] for all HTAN patients. Two CRLM patients (HT112C1-T1 and HT225C1) have multiple slices processed from a 3D tissue cube. We aligned the adjacent slices using PASTE2 [43] and provided the alignment matrix *W* to CalicoST infer CNAs and tumor clones in 3D space.

We used CalicoST to infer CNAs and tumor clones jointly across all five 10x Genomics Visium slices of the prostate organ. We also jointly identified germline heterozygous SNPs across the slices. Because there is no matched single-cell gene expression measurement, we treated all spots as pure in CalicoST. Since the slices are more likely to contain distinct clones because of their distant spatial location in the prostate organ, we initialized with five clones in CalicoST, which is higher than the default.

### 4.9 Evaluating the accuracy of inferred (allele-specific) copy numbers

We applied three metrics to evaluate the inferred allele-specific integer copy numbers: exact match, precision, and recall. Given *G* genome segments, the inferred allele-specific copy numbers 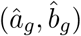 at bin *g*, and the ground truth allele-specific copy numbers (*a*_*g*_, *b*_*g*_), the exact match is the proportion of genomic segments where the inferred allele-specific copy numbers match the ground truth:

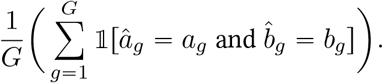

The precision is the proportion of predicted genome segments with CNAs that are supported by the ground truth, where a change of copy number in either A copy or B copy indicates the existence of CNA:

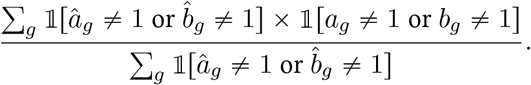

The recall is the proportion of genome segments with CNAs that are predicted:

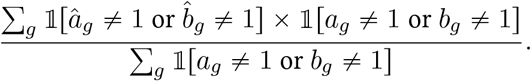

We extended the exact match to evaluate inferred copy number states (e.g. amplification state, deletion state) without integer copy numbers. With an abuse of notation, we denote 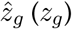 as the inferred (ground truth) copy number states at bin *g*. The exact match of copy number states is

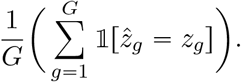

Numbat predicts 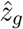 to be one of six copy number states: imbalanced amplification (amp), balanced amplification (bamp), balanced copy number neutral (neu), copy number neutral loss of heterozygosity (cnloh), imbalanced deletion (del), and balanced deletion (bdel). STARCH predicts 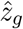 to be one of the three copy number states: amplification (amp), neutral (neu), and deletion (del). We converted the allele-specific integer copy numbers inferred by CalicoST or from WES to these states and compared the exact match values with the other methods.

### 4.10 Evaluating spatial coherence of tumor clones by z score of joincount statistics

We use joincount statistics [51, Chapter 3] to evaluate the spatial coherence of each inferred cancer clone. Joincount statistics describes the spatial autocorrelation of binary data. Given a weighted graph *G* = (*V, E, W*) where *W* is the weighted adjacency matrix, and let 𝓁 ∈ {0, 1}^|*V*|^ be the vertex label, the join-count statistics is the number of edges for which the two endpoints have labels {*a, b*}:

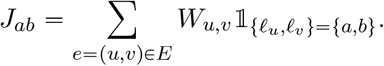

The z-score of joincount describes whether the number of edges is larger or smaller than the expectation assuming labels of the endpoints of each edge are i.i.d. samples from a Bernoulli distribution.

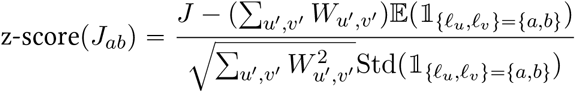

The higher the z-score of *J*_*ab*_ for *a* = *b*, the more spatially coherent the data is. Let *P*_*a*_ and *P*_*b*_ be the probability of *a* and *b* (*a, b* ∈ {0, 1}) in the Bernoulli distribution. The expectation and standard deviation is given by

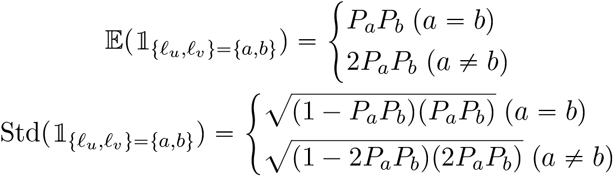

When there are multiple tumor clones, we compute the z-score of joincount, z-scorep*J*_11_q, for each clone by binarizing clone labels into whether each spot is in the given clone.

## Data Availability

Sequencing data are part of Human Tumor Atlas Network (HTAN) dbGaP Study Accession: phs002371.v3.p1 (https://www.ncbi.nlm.nih.gov/projects/gap/cgi-bin/study.cgi?study_id=phs002371.v3.p1), which will be released after publication. Sequencing data of the prostate cancer was obtained from the European Genome-phenome Archive (EGA) with accession EGAS00001006124.

## Code Availability

The code is publicly available at https://github.com/raphael-group/CalicoST under BSD 3-Clause license.

## Acknowledgements

This research is supported by NIH/NCI grants U24CA248453 and U24CA264027 to B.J.R and NIH grants U2CCA233303, U54AG075934, U24CA210972, R01HG009711 and R01CA260112 to L.D. C.M. is a Damon Runyon Fellow supported by the Damon Runyon Cancer Research Foundation (DRQ-15-22). We thank Palash Sashittal for his help with using Startle to reconstruct tumor phylogeny, and Uyen Mai for her help with inferring ancestor locations in phylogeography.

## Author Contributions

C.M and B.J.R conceived and designed the method. C.M. and B.J.R. developed the method. C.L. S.C. and L.D. curated and assisted with interpretation of the HTAN data. C.M. and M.B. performed the data analysis. C.M. M.B. and B.J.R wrote the manuscript. All authors read and approved the manuscript.

## References

[1] Marco Gerlinger, Andrew J Rowan, Stuart Horswell, James Larkin, David Endesfelder, Eva Gronroos, Pierre Martinez, Nicholas Matthews, Aengus Stewart, Patrick Tarpey, et al. Intratumor heterogeneity and branched evolution revealed by multiregion sequencing. New England journal of medicine, 366(10):883–892, 2012.

[2] Alexander M Frankell, Michelle Dietzen, Maise Al Bakir, Emilia L Lim, Takahiro Karasaki, Sophia Ward, Selvaraju Veeriah, Emma Colliver, Ariana Huebner, Abigail Bunkum, et al. The evolution of lung cancer and impact of subclonal selection in tracerx. Nature, pages 1–9, 2023.

[3] Emma Laks, Andrew McPherson, Hans Zahn, Daniel Lai, Adi Steif, Jazmine Brimhall, Justina Biele, Beixi Wang, Tehmina Masud, Jerome Ting, et al. Clonal decomposition and dna replication states defined by scaled single-cell genome sequencing. Cell, 179(5):1207–1221, 2019.

[4] Simone Zaccaria and Benjamin J Raphael. Characterizing allele-and haplotype-specific copy numbers in single cells with CHISEL. Nature Biotechnology, 39(2):207–214, 2021.

[5] Darlan C. Minussi, Michael D. Nicholson, Hanghui Ye, Alexander Davis, Kaile Wang, Toby Baker, Maxime Tarabichi, Emi Sei, Haowei Du, Mashiat Rabbani, Cheng Peng, Min Hu, Shanshan Bai, Yuwei Lin, Aislyn Schalck, Asha Multani, Jin Ma, Thomas O. McDonald, Anna Casasent, Angelica Barrera, Hui Chen, Bora Lim, Banu Arun, Funda Meric-Bernstam, Peter Van Loo, Franziska Michor, and Nicholas E. Navin. Breast tumours maintain a reservoir of subclonal diversity during expansion. Nature, 592(7853):302–308, apr 2021.

[6] Russell Schwartz and Alejandro A Schäffer. The evolution of tumour phylogenetics: principles and practice. Nature Reviews Genetics, 18(4):213–229, 2017.

[7] Maxime Tarabichi, Adriana Salcedo, Amit G Deshwar, Máire Ni Leathlobhair, Jeff Wintersinger, David C Wedge, Peter Van Loo, Quaid D Morris, and Paul C Boutros. A practical guide to cancer subclonal reconstruction from dna sequencing. Nature methods, 18(2):144–155, 2021.

[8] Moritz Gerstung, Clemency Jolly, Ignaty Leshchiner, Stefan C Dentro, Santiago Gonzalez, Daniel Rosebrock, Thomas J Mitchell, Yulia Rubanova, Pavana Anur, Kaixian Yu, et al. The evolutionary history of 2,658 cancers. Nature, 578(7793):122–128, 2020.

[9] Robert Noble, Dominik Burri, Cécile Le Sueur, Jeanne Lemant, Yannick Viossat, Jakob Nikolas Kather, and Niko Beerenwinkel. Spatial structure governs the mode of tumour evolution. Nature Ecology & Evolution, 6(2):207–217, 2022.

[10] Tongtong Zhao, Zachary D Chiang, Julia W Morriss, Lindsay M LaFave, Evan M Murray, Isabella Del Priore, Kevin Meli, Caleb A Lareau, Naeem M Nadaf, Jilong Li, et al. Spatial genomics enables multi-modal study of clonal heterogeneity in tissues. Nature, 601(7891):85–91, 2022.

[11] Andrew L Ji, Adam J Rubin, Kim Thrane, Sizun Jiang, David L Reynolds, Robin M Meyers, Margaret G Guo, Benson M George, Annelie Mollbrink, Joseph Bergenstråhle, et al. Multimodal analysis of composition and spatial architecture in human squamous cell carcinoma. Cell, 182(2):497–514, 2020.

[12] Dalia Barkley, Reuben Moncada, Maayan Pour, Deborah A Liberman, Ian Dryg, Gregor Werba, Wei Wang, Maayan Baron, Anjali Rao, Bo Xia, et al. Cancer cell states recur across tumor types and form specific interactions with the tumor microenvironment. Nature Genetics, 54(8):1192–1201, 2022.

[13] Jana Biermann, Johannes C Melms, Amit Dipak Amin, Yiping Wang, Lindsay A Caprio, Alcida Karz, Somnath Tagore, Irving Barrera, Miguel A Ibarra-Arellano, Massimo Andreatta, et al. Dissecting the treatment-naive ecosystem of human melanoma brain metastasis. Cell, 185(14):2591–2608, 2022.

[14] Yiping Wang, Joy Linyue Fan, Johannes C Melms, Amit Dipak Amin, Yohanna Georgis, Irving Barrera, Patricia Ho, Somnath Tagore, Gabriel Abril-Rodríguez, Siyu He, et al. Multimodal single-cell and whole-genome sequencing of small, frozen clinical specimens. Nature Genetics, 55(1):19–25, 2023.

[15] Anoop P Patel, Itay Tirosh, John J Trombetta, Alex K Shalek, Shawn M Gillespie, Hiroaki Wakimoto, Daniel P Cahill, Brian V Nahed, William T Curry, Robert L Martuza, et al. Single-cell RNA-seq highlights intratumoral heterogeneity in primary glioblastoma. Science, 344(6190):1396–1401, 2014.

[16] Ruli Gao, Shanshan Bai, Ying C Henderson, Yiyun Lin, Aislyn Schalck, Yun Yan, Tapsi Kumar, Min Hu, Emi Sei, Alexander Davis, et al. Delineating copy number and clonal substructure in human tumors from single-cell transcriptomes. Nature Biotechnology, 39(5):599–608, 2021.

[17] Rebecca Elyanow, Ron Zeira, Max Land, and Benjamin J Raphael. STARCH: Copy number and clone inference from spatial transcriptomics data. Physical Biology, 18(3):035001, 2021.

[18] Kieran R Campbell, Adi Steif, Emma Laks, Hans Zahn, Daniel Lai, Andrew McPherson, Hossein Farahani, Farhia Kabeer, Ciara O’Flanagan, Justina Biele, et al. clonealign: statistical integration of independent single-cell rna and dna sequencing data from human cancers. Genome Biology, 20(1):1–12, 2019.

[19] Pedro F Ferreira, Jack Kuipers, and Niko Beerenwinkel. Mapping single-cell transcriptomes to copy number evolutionary trees. In International Conference on Research in Computational Molecular Biology, pages 380–381. Springer, 2022.

[20] 10x Genomics. Spatial transcriptomics, 2021.

[21] Robert R Stickels, Evan Murray, Pawan Kumar, Jilong Li, Jamie L Marshall, Daniela J Di Bella, Paola Arlotta, Evan Z Macosko, and Fei Chen. Highly sensitive spatial transcriptomics at near-cellular resolution with Slide-seqV2. Nature biotechnology, 39(3):313–319, 2021.

[22] Jacqueline A Langdon, Jayne M Lamont, Debbie K Scott, Sara Dyer, Emma Prebble, Nick Bown, Richard G Grundy, David W Ellison, and Steven C Clifford. Combined genome-wide allelotyping and copy number analysis identify frequent genetic losses without copy number reduction in medulloblastoma. Genes, Chromosomes and Cancer, 45(1):47–60, 2006.

[23] Daisuke Kuga, Masahiro Mizoguchi, Yanlei Guan, Nobuhiro Hata, Koji Yoshimoto, Tadahisa Shono, Satoshi O Suzuki, Yoji Kukita, Tomoko Tahira, Shinji Nagata, et al. Prevalence of copy-number neutral loh in glioblastomas revealed by genomewide analysis of laser-microdissected tissues. Neuro-Oncology, 10(6):995–1003, 2008.

[24] Christine O’Keefe, Michael A McDevitt, and Jaroslaw P Maciejewski. Copy neutral loss of heterozygosity: a novel chromosomal lesion in myeloid malignancies. Blood, The Journal of the American Society of Hematology, 115(14):2731–2739, 2010.

[25] Mariam Jamal-Hanjani, Gareth A Wilson, Nicholas McGranahan, Nicolai J Birkbak, Thomas BK Watkins, Selvaraju Veeriah, Seema Shafi, Diana H Johnson, Richard Mitter, Rachel Rosenthal, et al. Tracking the evolution of non–small-cell lung cancer. New England Journal of Medicine, 376(22):2109–2121, 2017.

[26] Jean Fan, Hae-Ock Lee, Soohyun Lee, Da-eun Ryu, Semin Lee, Catherine Xue, Seok Jin Kim, Kihyun Kim, Nikolaos Barkas, Peter J Park, et al. Linking transcriptional and genetic tumor heterogeneity through allele analysis of single-cell RNA-seq data. Genome Research, 28(8):1217–1227, 2018.

[27] Teng Gao, Ruslan Soldatov, Hirak Sarkar, Adam Kurkiewicz, Evan Biederstedt, Po-Ru Loh, and Peter V Kharchenko. Haplotype-aware analysis of somatic copy number variations from single-cell transcriptomes. Nature Biotechnology, pages 1–10, 2022.

[28] Chi-Yun Wu, Anuja Sathe, Jiazhen Rong, Paul R Hess, Billy Lau, Susan M Grimes, Hanlee P Ji, and Nancy R Zhang. Cancer subclone detection based on DNA copy number in single cell and spatial omic sequencing data. bioRxiv, 2022.

[29] Chia-Kuei Mo, Jingxian Liu, Siqi Chen, Erik Storrs, Andre Luiz Targino da Costa, Michael D. Iglesia, Cong Ma, Reyka G. Jayasinghe, Andrew Houston, John M. Herndon, Jacqueline Mudd, Xinhao Liu, Alla Karpova, Andrew Shinkle, Austin N. Southard-Smith, Michael C. Wendl, S. Peter Goedegebuure, Abdurrahman Taha Mousa Ali Abdelzaher, Peng Bo, Lauren Fulghum, Samantha Livingston, Metin Balaban, Angela Hill, Joseph E. Ippolito, Vesteinn Thorsson, Jason M. Held, Eric H. Kim, Peter O. Bayguinov, Albert H. Kim, Kooresh I. Shoghi, Sidharth V. Puram, Tao Ju, Melissa A. Reimers, Cody Weimholt, Liang-I Kang, Deborah J. Veis, Milan G. Chheda, Russell Pachynski, Katherine C. Fuh, William E. Gillanders, Ryan C. Fields, Benjamin J. Raphael, Feng Chen, and Li Ding. Spatial clonal evolution and clone-specific microenvironment interactions within three-dimensional tumor structures. Nature, 2023. Submitted.

[30] Po-Ru Loh, Petr Danecek, Pier Francesco Palamara, Christian Fuchsberger, Yakir A Reshef, Hilary K Finucane, Sebastian Schoenherr, Lukas Forer, Shane McCarthy, Goncalo R. Abecasis, Richard Durbin, and Alkes L Price. Reference-based phasing using the Haplotype Reference Consortium panel. Nature Genetics, 48(11):1443–1448, nov 2016.

[31] Dylan M Cable, Evan Murray, Luli S Zou, Aleksandrina Goeva, Evan Z Macosko, Fei Chen, and Rafael A Irizarry. Robust decomposition of cell type mixtures in spatial transcriptomics. Nature Biotechnology, 40(4):517–526, 2022.

[32] Ying Ma and Xiang Zhou. Spatially informed cell-type deconvolution for spatial transcriptomics. Nature Biotechnology, pages 1–11, 2022.

[33] Simone Zaccaria, Mohammed El-Kebir, Gunnar W. Klau, and Benjamin J. Raphael. Phylogenetic Copy-Number Factorization of Multiple Tumor Samples. Journal of Computational Biology, 25(7):689–708, jul 2018.

[34] Roland F. Schwarz, Anne Trinh, Botond Sipos, James D. Brenton, Nick Goldman, and Florian Markowetz. Phylogenetic quantification of intra-tumour heterogeneity. PLOS Computational Biology, 10(4):1–11, 04 2014.

[35] Palash Sashittal, Henri Schmidt, Michelle M Chan, and Benjamin J Raphael. Startle: a star homoplasy approach for crispr-cas9 lineage tracing. bioRxiv, pages 2022–12, 2022.

[36] Simone Zaccaria and Benjamin J Raphael. Accurate quantification of copy-number aberrations and whole-genome duplications in multi-sample tumor sequencing data. Nature Communications, 11(1):1–13, 2020.

[37] Susanne Schulze and Iver Petersen. Gender and ploidy in cancer survival. Cellular Oncology, 34:199–208, 2011.

[38] Olagunju A Ogunbiyi, Paul J Goodfellow, Klaus Herfarth, Guiseppe Gagliardi, Paul E Swanson, Elisa H Birnbaum, Thomas E Read, James W Fleshman, Ira J Kodner, and Jeffrey F Moley. Confirmation that chromosome 18q allelic loss in colon cancer is a prognostic indicator. Journal of Clinical Oncology, 16(2):427–433, 1998.

[39] Jin Jen, Hoguen Kim, Steven Piantadosi, Zong-Fan Liu, Roy C Levitt, Pertti Sistonen, Kenneth W Kinzler, Bert Vogelstein, and Stanley R Hamilton. Allelic loss of chromosome 18q and prognosis in colorectal cancer. New England Journal of Medicine, 331(4):213–221, 1994.

[40] L1 Cawkwell, FA Lewis, and P Quirke. Frequency of allele loss of dcc, p53, rbi, wt1, nf1, nm23 and apc/mcc in colorectal cancer assayed by fluorescent multiplex polymerase chain reaction. British Journal of Cancer, 70(5):813–818, 1994.

[41] Keizou Ookawa, Michiie Sakamoto, Setsuo Hirohashi, Yutaka Yoshida, Takashi Sugimura, Masaaki Terada, and Jun Yokota. Concordant p53 and dcc alterations and allelic losses on chromosomes 13q and 14q associated with liver metastases of colorectal carcinoma. International journal of cancer, 53(3):382–387, 1993.

[42] Anoop P. Patel, Itay Tirosh, John J. Trombetta, Alex K. Shalek, Shawn M. Gillespie, Hiroaki Wakimoto, Daniel P. Cahill, Brian V. Nahed, William T. Curry, Robert L. Martuza, David N. Louis, Orit Rozenblatt-Rosen, Mario L. Suvàz, Aviv Regev, and Bradley E. Bernstein. Single-cell RNA-seq highlights intratumoral heterogeneity in primary glioblastoma. Science, 344(6190):1396–1401, jun 2014.

[43] Xinhao Liu, Ron Zeira, and Benjamin J Raphael. Partial alignment of multislice spatially resolved transcriptomics data. Genome Research, 2023.

[44] Andrew Erickson, Mengxiao He, Emelie Berglund, Maja Marklund, Reza Mirzazadeh, Niklas Schultz, Linda Kvastad, Alma Andersson, Ludvig Bergenstråhle, Joseph Bergenstråhle, et al. Spatially resolved clonal copy number alterations in benign and malignant tissue. Nature, 608(7922):360–367, 2022.

[45] Alexander T El Gammal, Michael Brüchmann, Jozef Zustin, Hendrik Isbarn, Olaf JC Hellwinkel, Jens Köllermann, Guido Sauter, Ronald Simon, Waldemar Wilczak, Jörg Schwarz, et al. Chromosome 8p deletions and 8q gains are associated with tumor progression and poor prognosis in prostate cancer. Clinical Cancer Research, 16(1):56–64, 2010.

[46] Paul CMS Verhagen, Karin GL Hermans, Mariel O Brok, Wytzke M van Weerden, Marcel GJ Tilanus, Roel A de Weger, Tom A Boon, and Jan Trapman. Deletion of chromosomal region 6q14-16 in prostate cancer. International Journal of Cancer, 102(2):142–147, 2002.

[47] Martina Kluth, Simon Jung, Omar Habib, Mina Eshagzaiy, Anna Heinl, Nina Amschler, Sawinee Masser, Malte Mader, Frederic Runte, Philipp Barow, et al. Deletion lengthening at chromosomes 6q and 16q targets multiple tumor suppressor genes and is associated with an increasingly poor prognosis in prostate cancer. Oncotarget, 8(65):108923, 2017.

[48] Chi-Yun Wu, Billy T Lau, Heon Seok Kim, Anuja Sathe, Susan M Grimes, Hanlee P Ji, and Nancy R Zhang. Integrative single-cell analysis of allele-specific copy number alterations and chromatin accessibility in cancer. Nature Biotechnology, 39(10):1259–1269, 2021.

[49] Julien Stoehr. A review on statistical inference methods for discrete markov random fields. arXiv preprint 1704.03331, 2017.

[50] Julian Besag. Spatial interaction and the statistical analysis of lattice systems. Journal of the Royal Statistical Society: Series B (Methodological), 36(2):192–225, 1974.

[51] BOUAYAD AGHA Salima and MD Bellefon. Spatial autocorrelation indices, pages 60–62. 2018.

[52] Xianjie Huang and Yuanhua Huang. Cellsnp-lite: an efficient tool for genotyping single cells. Bioinformatics, 37(23):4569–4571, 2021.

[53] Yuchao Jiang, Yu Qiu, Andy J Minn, and Nancy R Zhang. Assessing intratumor heterogeneity and tracking longitudinal and spatial clonal evolutionary history by next-generation sequencing. Proceedings of the National Academy of Sciences, 113(37):E5528–E5537, 2016.

