## Supplementary notes and figures for "Inferring allele-specific copy number aberrations and tumor phylogeography from spatially resolved transcriptomics"

### Supplementary Information

#### S1 Extracting allele and total counts

We follow the pipeline of Numbat [27] to extract allele and total counts. Using the 1000 Genomes SNP panel, we genotype the reference allele (or A allele) and alternative allele (or B allele) using cellsnp-lite [52]. The output contains the B allele counts and total counts summed across all spots, and the counts of individual spots at each SNP loci. We treat a SNP locus as homozygous if the frequency of B alleles when summed across all spots is close to 0 or 1 by a threshold. All homozygous and heterozygous SNPs are retained for reference-based phasing using Eagle2 [30], but homozygous SNPs are removed from the B allele counts $\tilde{\mathbf{Y}}$ and total allele counts $\tilde{\mathbf{D}}$.

#### S2 Copy number probabilistic model for the heterogeneous clones case

Suppose each spot $n$ is a mixture of $M$ clones and a proportion of $\tilde{\theta}_{m,n}$ of the counts are from clone $m$, $\sum_{m=1}^{M} \tilde{\theta}_{m,n} = 1$. We adapt the mean of the Negative Binomial distribution of $\mathbf{X}$ by a mixture of RDR values. With a little abuse of notation, we denote $\mu_{g,m}$ as the RDR value that segment $g$ in clone $m$ takes among the $K$ unique values, i.e. $\sum_{k=1}^{K} \mathbb{1}[z_{g,m} = k]\mu_{k,m}$.

$$x_{g,n} \sim NB\left(T_n \lambda_g \sum_m \tilde{\theta}_{m,n} \mu_{m,g}, \phi\right). \tag{11}$$

The Beta-binomial distribution of $\mathbf{Y}$ under the mixture is parameterized by

$$y_{g,n} \sim BetaBinom\left(D_g, \tau \frac{\sum_m \tilde{\theta}_{m,n} \mu_{g,m} p_{g,m}}{\sum_m \tilde{\theta}_{m,n} \mu_{g,m}}, \tau(1 - \frac{\sum_m \tilde{\theta}_{m,n} \mu_{g,m} p_{g,m}}{\sum_m \tilde{\theta}_{m,n} \mu_{g,m}})\right). \tag{12}$$

With a similar change in notation of $\mu_{g,m}$ and $p_{g,m}$. See Section S8 for the derivation of (11) and (12).

However, deconvolving the input matrices into multiple cancer clones is challenging and the solutions are potentially non-identifiable. Therefore, we make the simplifying assumption that each spot may only contain at most one cancer clone but can be mixed with normal cells, that is $|\{m \in \text{cancer clones} : \tilde{\theta}_{m,n} > 0\}| \leqslant 1$. Because the RDR and BAF of normal cells are known values, $\mu_{g,m} = 1, p_{g,m} = 0.5$, inferring the RDR and BAF of the only cancer clone in a spot becomes identifiable. We also simplify the notation and define the *tumor count proportion* $\boldsymbol{\theta} = [\theta_n]$ as the proportion of counts from the only cancer clone of each spot if not purely normal, $\theta_n = \sum_{m \in \text{cancer clones}} \tilde{\theta}_{m,n}$. Let $\boldsymbol{\ell} = [\ell_n]$ be the label of the only cancer clone, we rewrite the probabilistic model of $\mathbf{X}$ and $\mathbf{Y}$ under the simplifying assumption:

$$x_{g,n} \mid \ell_n = m \sim NB\left(T_n \lambda_g (\theta_n \mu_{m,g} + 1 - \theta_n), \phi\right) \tag{13}$$

$$y_{g,n} \mid \ell_n = m \sim BetaBinom\left(D_g, \tau \frac{\theta_n \mu_{g,m} p_{g,m} + 0.5(1 - \theta_n)}{\theta_n \mu_{g,m} + 1 - \theta_n}, \tau(1 - \frac{\theta_n \mu_{g,m} p_{g,m} + 0.5(1 - \theta_n)}{\theta_n \mu_{g,m} + 1 - \theta_n})\right). \tag{14}$$

We take $\theta_n$ estimated from cell-type deconvolution methods (e.g., RCTD [31] and CARD [32]) as input assuming a reference of cell-type specific expression is available. Note that we define the proportion $\boldsymbol{\theta}$ as the proportion of counts, instead of the proportion of cells. We claim that the output from cell-type deconvolution methods matches our proportion definition. Different cell types may have different total transcript counts within each cell; by normalizing all cell types to have the same total transcript count in cell-type deconvolution, those methods infers the proportions of counts of each cell type rather than proportions of cells.

### S3 Phasing

We aggregate the allele counts from the same haplotype across adjacent SNP loci in $\tilde{\mathbf{Y}}$ to get $\mathbf{Y}$. However, a haplotype may either be the A allele or the B allele at different SNP loci. The assignment from A and B alleles to each haplotype is called *phasing*. Phasing is shared across all spots by definition, we seek to infer the phasing across all spots.

This phasing inference problem has been studied in previous methods for inferring CNAs in bulk WGS data, single-cell DNA-seq data, as well as scRNA-seq data [4, 27, 36]. Based on a reference-based phasing (e.g. Eagle2), these methods seek to correct the phase-switch errors made by reference-based phasing by modeling and maximizing the likelihood of observed BAF signals along the genome. Particularly, Numbat [27] seeks to infer CNA states and phasing together under one hidden Markov model with pre-defined 15 states. Observing the phasing is shared across all cells and spots, we simplified Numbat's model and used it only for phasing.

Let the phasing be $\mathbf{h} = [h_g] \in \{A, B\}^G$ for each SNP loci $g$. We model the B allele count after reference-based phasing using $h_g$ and an additional BAF cluster hidden state $z_g$ (with a little abuse of notation, we use the same variable $z_g$ here as with the copy number states). Then, the B allele count can be modeled by

$$\sum_n \tilde{y}_{g,n} \mid h_g = B; z_g = i \sim BetaBinom(\sum_n \tilde{d}_{g,n}, \tau p_i, \tau(1-p_i))$$
$$\sum_n \tilde{y}_{g,n} \mid h_g = A; z_g = i \sim BetaBinom(\sum_n \tilde{d}_{g,n}, \tau(1-p_i), \tau p_i),$$

which is the emission probability of the HMM. Note that B allele counts are conditioned on two hidden variables, phasing $h_g$ and BAF cluster $z_g$, and these two variables compose a composite hidden state of the overall HMM. We assume the two components of the composite state are independent in the start probability and state-transition probability:

$$\mathbb{P}((z_g, h_g) \mid (z_{g-1}, h_{g-1})) = \mathbb{P}(z_g \mid z_{g-1})\mathbb{P}(h_g \mid h_{g-1}).$$

The transition among BAF states and phase is defined respectively as follows

$$\mathbb{P}(z_g \mid z_{g-1}) = \begin{cases} t & (z_g = z_{g-1}) \\ \dfrac{1-t}{K-1} & (z_g \neq z_{g-1}). \end{cases}$$
$$\mathbb{P}(h_g \mid h_{g-1}) = \begin{cases} q & (h_g = h_{g-1}) \\ 1-q & (h_g \neq h_{g-1}). \end{cases}$$

We apply the HMM model to the pseudobulk of all spots and estimate using Baum-Welch algorithm, thus inferring one phase vector $\mathbf{h}$ across all spots. Numbat [27], on the other hand, infers an independent phase along with the CNA states for each cancer clone, potentially inducing errors in both phasing and CNA inference.

### S4 Identifying a confident set of normal spots and obtaining baseline transcript counts $\boldsymbol{\lambda}$

If the tumor count proportions $\theta_n$ are available, we select the set of spots for which $\theta_n$ is below a threshold as confident normal spots. The threshold is chosen such that the transcript counts of each genome segment on average is at least 200 when aggregated across confident normal spots.

If the tumor count proportions are unavailable, CalicoST identifies a confident set of normal spots with the closest-to-balanced BAF along the genome. It is challenging to evaluate for individual spots whether the BAF is close to balance because total allele counts are sparse. Therefore, we group spots with similar BAF signals along the genome and evaluate the BAF of each group. We infer the grouping by solving the **CNA and clone inference problem** using only the allele count matrices $\mathbf{Y}, \mathbf{D}$. Let $\mathbf{p}^0 \in \mathbb{R}^K$ be the estimated BAF across $K$ states, and $\mathbf{Z}^0 = [z^0_{g,m}]$ be the state indicator for each genome segment in each group. Hence, normal spots must be within the group $m'$ with the most balanced BAFs $m' = \arg\min_m \sum_{g=1}^{G} |\sum_{k=1}^{K} \mathbb{1}[z_{g,m} = k]p^0_k - 0.5|$. To avoid the case where a small cancer clones are mixed within spot group $m'$, we select a given number of spots with the smallest variance of log-transformed transcript counts along genome inside clone $m'$ as confident normal spots.

Let $J$ denote the set of confident normal spots, the transcript count proportions across genome segments in normal cells $\boldsymbol{\lambda}$ are derived by averaging the expression across confident normal spots $J$ and normalizing to sum to 1, $\lambda_g \propto \sum_{i \in J} x_{g,i}$.

#### S5 Filtering genes and segments based on identified normal spots

Allele imbalance occasionally can be caused by allele-specific expression, e.g. cis-regulatory elements, and we seek to identify and filter out genomic segments that are potentially affected by allele-specific expression. We assume allele-specific expression affects normal cells and the B counts of affected genomic segments cannot be well explained by BAF 0.5. Let $J$ be the confident set of normal cells and let $F_{\text{Betabinom}}(y; d, \tau p, \tau(1-p))$ be the CDF of a Beta-binomial distribution with $d$ total allele counts and $p$ B allele frequency. The set of genomic segments that are affected by allele-specific expression is

$$\{g \in G : F_{\text{Betabinom}}(\sum_{i \in J} y_{g,i}; \sum_{i \in J} d_{g,i}, 0.5\tau, 0.5\tau) < \text{threshold or}$$
$$F_{\text{Betabinom}}(\sum_{i \in J} y_{g,i}; \sum_{i \in J} d_{g,i}, 0.5\tau, 0.5\tau) > 1 - \text{threshold}\}$$

We filtered out these genomic segments from $\mathbf{Y}$ and $\mathbf{D}$.

Differential expressions, though usually caused by other biological factors than CNAs, can dominate the estimation of RDR $\boldsymbol{\mu}$ when the transcript count of the corresponding gene is high. To reduce the effects from other biological factors in estimating $\boldsymbol{\mu}$, we filter out differentially expressed genes between confident normal spots and tumor ones, and between confident normal spots and other spots in initial clone $m'$ if (1) their total transcript counts across all spots are above the 80% quantile and (2) the log fold change of expression is above a threshold. We argue that CalicoST can still accurately estimate $\boldsymbol{\mu}$ after filtering the above set of genes. Because large CNAs alter the expression of many adjacent genes along genome, the remaining genes can still indicate copy number gains or a losses.

#### S6 Inferring allele-specific integer copy numbers per clone

CalicoST estimates the integer allele-specific copy numbers $\mathbf{A}$ and $\mathbf{B}$ using the estimated RDR $\boldsymbol{\mu}$, BAF $\mathbf{p}$, and copy number states $\mathbf{Z}$. Because the RDR and BAF correspond to the transformation from $\mathbf{A}$ and $\mathbf{B}$ in equation (5), we optimize over $\mathbf{A}$ and $\mathbf{B}$ to minimize the deviation between the transformed RDR and BAF values and the estimated ones in (2).

Since $\mathbf{Z}$ groups the genome segments by copy number states, we only estimate a single allele-specific copy number for the state within each clone $m$. We introduce the notation $\tilde{\mathbf{A}} = [\tilde{a}_{k,m}]$ for the $K$ values of A copy and $\tilde{\mathbf{B}} = [\tilde{b}_{k,m}]$ for the $K$ values of B copy for each clone $m$. Given the copy number states $\mathbf{Z}$, the allele-specific copy numbers $\mathbf{A}, \mathbf{B}$ along the entire genome can be expressed using $\tilde{\mathbf{A}}, \tilde{\mathbf{B}}$:

$$a_{g,m} = \sum_{k=1}^{K} \mathbb{1}_{z_{g,m}=k} \tilde{a}_{k,m}.$$

In addition, the transformation to RDR and BAF from $\tilde{\mathbf{A}}, \tilde{\mathbf{B}}$ is

$$\mu_k = \frac{\tilde{a}_{k,m} + \tilde{b}_{g,m}}{\sum_{k=1}^{K} \tilde{\lambda}_{k,m}(\tilde{a}_{k,m} + \tilde{b}_{k,m})}$$
$$p_k = \frac{\tilde{b}_{k,m}}{\tilde{a}_{k,m} + \tilde{b}_{k,m}},$$

where $\tilde{\lambda}_{k,m} = \sum_g \mathbb{1}_{z_{g,m}=k} \lambda_g$.

Generally, the integer copy numbers are not identifiable; different $\tilde{\mathbf{A}}, \tilde{\mathbf{B}}$ may lead to the same approximation of $\boldsymbol{\mu}, \mathbf{p}$, for example, scaling both $\tilde{\mathbf{A}}$ and $\tilde{\mathbf{B}}$ by the same integer multiplier. Therefore, we constrain the allele-specific integer copy numbers not to be too large and the ploidy of the genome $\psi_m$ not to be too high. Denoting the constrained space as $\mathcal{C}$, we solve the following constrained optimization problem for each clone $m$ to infer $\tilde{\mathbf{A}}_{\cdot,m}$ and $\tilde{\mathbf{B}}_{\cdot,m}$:

$$\begin{aligned} \underset{(\psi_m, \tilde{\mathbf{A}}_{\cdot,m}, \tilde{\mathbf{B}}_{\cdot,m}) \in \mathcal{C}}{\arg\min} \quad & \left( \sum_{k=1}^{K} \left( w_k^{\text{RDR}} |\mu_k - \frac{\tilde{a}_{k,m} + \tilde{b}_{g,m}}{\psi_m}|^2 + w_k^{\text{BAF}} |p_k - \frac{\tilde{b}_{k,m}}{\tilde{a}_{k,m} + \tilde{b}_{k,m}}|^2 \right) \right) + \\ & w^{\text{ploidy}} \|\psi_m - \sum_{k=1}^{K} \tilde{\lambda}_{k,m}(\tilde{a}_{k,m} + \tilde{b}_{k,m})\|_2^2, \end{aligned} \tag{15}$$

where $w_k^{\text{RDR}}$, $w_k^{\text{BAF}}$ and $w^{\text{ploidy}}$ are weights.

Different weights lead to different optimum in allele-specific integer copy numbers. In CalicoST, we implemented two settings of the weights and constraints. In the first setting, we set $w^{\text{ploidy}} = \infty$, $w_k^{\text{RDR}} = 0.3 w_k^{\text{BAF}} = \sum_{g=1}^{G} \mathbb{1}[z_{g,m} = k]$, where we applies a scaling of $w_k^{\text{RDR}}$ to account for the different dynamic range of RDR (ranging from 0 to an integer total copy number divided by 2) and BAF (ranging from 0 to 1). We set the constrained space $\mathcal{C}$ as $\mathcal{C}$ as $\mathcal{C} = \{\tilde{a}_{k,m} \leqslant 4 (\forall k), \tilde{b}_{k,m} \leqslant 4 (\forall k), \psi_m < 3\}$.

In the second setting, we borrow the idea from CHISEL [36] and identify a balanced diploid state $j$ such that estimated BAF $p_j$ is close to 0.5 and estimated RDR $\mu_j$ is the lowest. The balanced diploid state will have integer copy numbers $\tilde{a}_{j,m} = \tilde{b}_{j,m} = 1$. Using the balanced diploid state, we set the constrained space as $\mathcal{C}$ as $\mathcal{C}$ as $\mathcal{C} = \{\tilde{a}_{j,m} = \tilde{b}_{j,m} = 1, \tilde{a}_{k,m} \leqslant 4 (\forall k), \tilde{b}_{k,m} \leqslant 4 (\forall k), \psi_m < 3\}$, and set $w^{\text{ploidy}} = 0$, $w_j^{\text{RDR}} = \infty$, and $w_k^{\text{RDR}} = 0.3 w_k^{\text{BAF}} = \sum_{g=1}^{G} \mathbb{1}[z_{g,m} = k]$ for the remaining $k$ as above.

Note that CHISEL's objective function for estimating integer copy numbers is a slight variation of (15). In CHISEL, $w^{\text{ploidy}}$ is set to 0 as our second setting. CHISEL models observed RDR and BAF using Gaussian distributions and uses the inverse of Gaussian variance as $w_k^{\text{BAF}}$ and $w_k^{\text{RDR}}$. In addition, instead of constraining to space $\mathcal{C}$, CHISEL incorporates a penalty term of $\psi_m$. Because the gene expression in SRT has higher variance than the read coverage in scDNA-seq, there may not be a robust penalty weight for $\psi_m$ across various SRT samples. Therefore, we enforce the constraint $\mathcal{C}$ in CalicoST instead of using penalty.

### S7 Choosing the number of clones

CalicoST requires an initial number of clones provided by users and uses a statistics to evaluate the similarity of CNA profiles and combine initial clones. Even though model selection criteria such as AIC and

BIC have been proposed, it is hard to directly apply them in our model, where an HMM model is nested in an HMRF. Instead, we evaluate the similarity between CNA profiles of two clones using the following statistics and merge them if their similarity passes a threshold.

Given two clones $m$ and $m'$ with their HMM states $\mathbf{Z}_{\cdot,m}$ and $\mathbf{Z}_{\cdot,m'}$, we suppose their HMM states differ in $I$ intervals of genomic bins, $R_1, \ldots, R_I$, where each interval has a consistent copy number state within each clone. Let $\mathbf{Z}_{R_i,m} = (z_{g,m})_{g \in R_i}$ and $\mathbf{Z}_{R_i,m'} = (z_{g,m'})_{g \in R_i}$, which are constant vectors by our construction of $R_i$. If the CNAs of clone $m$ and $m'$ are similar in interval $R_i$, the probabilities of observing the count data under copy number states $\mathbf{Z}_{R_i,m}$ should be similar to those under copy number states $\mathbf{Z}_{R_i,m'}$. We use the Neyman Pearson statistics to capture the ratio between the probabilities under the two copy number states:

$$T_{m,m',R_i} = \frac{\mathbb{P}(\mathbf{X}_{R_i,m}, \mathbf{Y}_{R_i,m} \mid \mathbf{Z}_{R_i,m}, \boldsymbol{\mu}, \mathbf{p})\mathbb{P}(\mathbf{X}_{R_i,m'}, \mathbf{Y}_{R_i,m'} \mid \mathbf{Z}_{R_i,m'}, \boldsymbol{\mu}, \mathbf{p})}{\mathbb{P}(\mathbf{X}_{R_i,m}, \mathbf{Y}_{R_i,m} \mid \mathbf{Z}_{R_i,m'}, \boldsymbol{\mu}, \mathbf{p})\mathbb{P}(\mathbf{X}_{R_i,m'}, \mathbf{Y}_{R_i,m'} \mid \mathbf{Z}_{R_i,m}, \boldsymbol{\mu}, \mathbf{p})}.$$

$T_{m,m',R_i} \approx 1$ indicates the probabilities are indeed similar and the CNAs of clone $m$ in interval $R_i$ are similar to those of clone $m'$; while $T_{m,m',R_i} \gg 1$ indicates CNAs of clone $m$ largely differ from those of clone $m'$ in interval $R_i$. CalicoST decides to merge a set of initial clones if the CNAs of all pairs of clones are similar across all intervals, $T_{m,m',R_i} \leqslant 1 + \epsilon$ for all clone pairs $m, m'$ for all intervals $R_i$ under a user-defined threshold $\epsilon$.

### S8 Derivation of the probabilistic model for the heterogeneous clone case

#### S8.1 Negative Binomial model for $\mathbf{X}$

We use a Negative Binomial distribution to empirically approximate a Dirichlet-Multinomial distribution for expression counts. We start with deriving the Dirichlet-Multinomial distribution first. We use the following notations defined previously: total UMI counts $T_n$, the probability $\boldsymbol{\lambda}$ of sequencing a UMI in normal cells, the proportion $\tilde{\theta}_{m,n}$ of UMIs from clone $m$, A and B allele copy numbers per bin per clone $\mathbf{A} = [a_{g,m}], \mathbf{B} = [b_{g,m}] \in \mathbb{Z}^{G \times M}$. Let the probability of sequencing a UMI at each bin in cell mixture of be $\boldsymbol{\alpha} \in \mathbb{R}^G$, which is the parameter in the Dirichlet-Multinomial distribution $DirMult(T, \boldsymbol{\alpha})$.

$$\begin{aligned}\alpha_g &\propto \sum_m \theta_{m,n} \mathbb{P}(\text{sequencing a UMI from bin } g \mid \text{clone } m) \\ &= \sum_m \theta_{m,n} \frac{\lambda_g (a_{g,m} + b_{g,m})}{\sum_{g'} \lambda_{g'} (a_{g',m} + b_{g',m})}.\end{aligned}$$

Using the definition of RDR $\mu_{g,m} = \frac{a_{g,m}+b_{g,m}}{\sum_g \lambda_g (a_{g,m}+b_{g,m})}$, we simplify $\boldsymbol{\alpha}$ by

$$\alpha_g \propto \sum_m \theta_m \lambda_g \mu_{g,m}.$$

$\alpha_g$ should satisfy $\sum_{g'} \alpha_{g'} = 1$, and we can verify that $\sum_g \sum_m \theta_m \lambda_g \mu_{g,m} = 1$. Therefore,

$$\alpha_g = \sum_m \theta_m \lambda_g \mu_{g,m}.$$

As a result, the Negative Binomial distribution that approximates the Dirichlet-Multinomial distribution is

$$NB(T\alpha_g, \phi) = NB(T\lambda_g \sum_m \theta_m \mu_{g,m}, \phi).$$

#### S8.2 Beta-binomial model for $\mathbf{Y}$

In a mixture of clones with UMI proportion per clone $\theta_{m,n}$ and clone-specific BAF $p_{g,m}$, the mixture BAF in the Beta-binomial distribution (11) is

$$\begin{aligned}
\text{mixture BAF} &= \frac{\sum_m \theta_{m,n}\mathbb{P}(\text{sequencing a UMI from bin } g)p_{g,m}}{\sum_m \theta_m \mathbb{P}(\text{sequencing a UMI from bin } g)} \\
&= \frac{\sum_m \theta_{m,n}\frac{\lambda_g(a_{g,m}+b_{g,m})}{\sum_{g'}\lambda_{g'}(a_{g',m}+b_{g',m})}p_{g,m}}{\sum_m \theta_{m,n}\frac{\lambda_g(a_{g,m}+b_{g,m})}{\sum_{g'}\lambda_{g'}(a_{g',m}+b_{g',m})}} \\
&= \frac{\sum_m \theta_{m,n}\lambda_g\mu_{g,m}p_{g,m}}{\sum_m \theta_{m,n}\lambda_g\mu_{g,m}} \\
&= \frac{\sum_m \theta_{m,n}\mu_{g,m}p_{g,m}}{\sum_m \theta_{m,n}\mu_{g,m}}
\end{aligned}$$

#### S8.3 Relationship between tumor UMI proportion $\theta_n$, tumor purity $\rho$, and cancer cell fraction (CCF)

Whole genome sequencing (WGS) technology measures DNA of mixed clones. Previous methods [7, 53] for studying single-nucleotide variants (SNVs) and CNAs on WGS data express RDR and BAF in the bulk mixture using the *tumor purity* and *cancer cell fraction (CCF)*. Tumor purity $\rho$ is the fraction of tumor cells in the bulk of cells. CCF is the fraction of tumor cells carrying a given somatic mutation. Under our assumption that the bulk contains one tumor clone besides normal cells, CCF=1. We denote A allele copy number as $\mathbf{A} \in \mathbb{Z}^G$ and B allele copy number $\mathbf{B} \in \mathbb{Z}^G$ for the tumor clone. And the normal clone has one A copy and one B copy for the entire genome. Previous work has the following expression of a BAF of the bulk of cells at bin $g$:

$$\text{bulk BAF} = \frac{\rho b_g + (1-\rho)}{\rho(a_g + b_g) + 2(1-\rho)}. \tag{16}$$

We use tumor UMI proportion $\theta_n$ to express RDR and BAF in the bulk mixture of SRT data in Section S2. Our BAF formula in equation (14) naturally extends to WGS. Specifically $\theta_n$ now represents the proportion of reads rather than UMIs from tumor cells, and RDR $\mu_g$ the tumor clone is a function of copy number of segment length $l_g$ of bin $g$ and total genome length $L = \sum_g l_g$, $\mu_g = \frac{a_g+b_g}{\sum_g l_g/L(a_g+b_g)}$. For simplicity, we drop the subscription that specifies spot $n$. Then our formula for bulk BAF extended to WGS is

$$\text{bulk BAF} = \frac{\theta\mu_g p_g + 0.5(1-\theta)}{\theta\mu_g + (1-\theta)}. \tag{17}$$

We will show that our formula and (16) are equivalent in the next.

**Theorem 1.** *The bulk BAF expressions in equation (16) and in equation (17) are equivalent in WGS data.*

The key to proving the equivalence is the relationship between tumor purity $\rho$ and tumor read proportion $\theta$. WGS data satisfies the following probability in generating sequencing reads, through which $\theta$ can be expressed by $\rho$.

**Property 1.** *Given tumor purity $\rho$, the normal genome that is partitioned into $G$ bins, the length $l_g$ of bin $g$, the probability of sequencing read $r$ coming from tumor (or normal) cells is proportional to the fraction of* *basepairs from tumor (or normal) genome.*

$$\mathbb{P}(r \in \textit{tumor}) \propto \rho \sum_{g=1}^{G} (a_g + b_g) l_g$$
$$\mathbb{P}(r \in \textit{normal}) \propto \rho \sum_{g=1}^{G} 2 l_g.$$

Therefore, the tumor read proportion $\theta$ is

$$\theta = \frac{\mathbb{P}(r \in \text{tumor})}{\mathbb{P}(r \in \text{tumor}) + \mathbb{P}(r \in \text{normal})} = \frac{\rho \sum_g (a_g + b_g) l_g}{\rho \sum_g (a_g + b_g) l_g + (1-\rho) \sum_g 2 l_g}$$

*Proof of Theorem 1.* Dividing the numerator and denominator of $\theta$ by $L$ gives $\theta = \frac{\frac{1}{L}\rho \sum_g (a_g+b_g) l_g}{\frac{1}{L}\rho \sum_g (a_g+b_g) l_g + \frac{1}{L}(1-\rho)\sum_g 2 l_g}$. Also let the denominator of $\theta$ be $\text{Denom} = \frac{1}{L}\rho \sum_g (a_g+b_g) l_g + \frac{1}{L}(1-\rho)\sum_g 2 l_g$. Using these notations, $\theta \mu_g$ is

$$\theta \mu_g = \frac{\frac{1}{L}\rho \sum_g (a_g + b_g) l_g}{\text{Denom}} \frac{(a_g + b_g)}{\frac{1}{L}\sum_g (a_g + b_g) l_g} = \frac{\rho (a_g + b_g)}{\text{Denom}}.$$

Plugging this formula of $\theta \mu_g$ and $1 - \theta = \frac{(1-\rho)\sum_g 2 l_g}{\text{Denom}}$ in equation (17), we get

$$\begin{aligned}
\text{bulk BAF} &= \frac{\rho (a_g + b_g) p_g + 0.5 \frac{1}{L}(1-\rho)\sum_g 2 l_g}{\rho (a_g + b_g) + \frac{1}{L}(1-\rho)\sum_g 2 l_g} \\
&= \frac{\rho (a_g + b_g) p_g + (1-\rho)}{\rho (a_g + b_g) + 2(1-\rho)} \\
&= \frac{\rho (a_g + b_g) \frac{b_g}{a_g + b_g} + (1-\rho)}{\rho (a_g + b_g) + 2(1-\rho)} \\
&= \frac{\rho b_g + (1-\rho)}{\rho (a_g + b_g) + 2(1-\rho)}
\end{aligned}$$

which is the same as equation (16). □

However, Property (1) does not hold in SRT data. SRT measures gene expression, and the total expression in tumor and normal cells is not proportional to their numbers of genomic base pairs. We cannot derive an expression of tumor UMI proportion using tumor purity. Therefore, we use bulk BAF formula (17) with tumor UMI proportion $\theta$.

#### S9 Evaluating likelihoods for pooled spots and individual spots

**Theorem 2.** *If we simplifying the distribution of* $\mathbf{X}$ *to Poisson distributions and* $\mathbf{Y}$ *to Binomial distributions,*

$$X_{g,n} \sim Pois(T_n \mu_g^m)$$
$$Y_{g,n} \sim B(D_{g,n}, p_g^m, (1 - p_g^m)),$$

*then the optima of HMM objective of individual spots are the same as that of pseudobulk.*

$$\underset{\boldsymbol{\mu}^m, \mathbf{p}^m}{\arg\max} \sum_{\mathbf{Z}_{\cdot,m}} \mathbb{P}(\mathbf{X}_{\cdot,I_m}, \mathbf{Y}_{\cdot,I_m}, \mathbf{D}_{\cdot,I_m} \mid \mathbf{Z}_{\cdot,m}, \boldsymbol{\mu}^m, \mathbf{p}^m))\mathbb{P}(\mathbf{Z}_{\cdot,m})$$

$$= \underset{\boldsymbol{\mu}^m, \mathbf{p}^m}{\arg\max} \sum_{\mathbf{Z}_{\cdot,m}} \mathbb{P}(\sum_{n\in I_m} \mathbf{X}_{\cdot,n}, \sum_{n\in I_m} \mathbf{Y}_{\cdot,n}, \sum_{n\in I_m} \mathbf{D}_{\cdot,n} \mid \mathbf{Z}_{\cdot,m}, \boldsymbol{\mu}^m, \mathbf{p}^m)\Big)\mathbb{P}(\mathbf{Z}_{\cdot,m})$$

*Proof.* We first show that the likelihoods of pooled and individual spots are different by a constant scalar. Without loss of generality, we assume $\mathbf{Z}_{g,m} = k$ at bin $g$ in clone $m$, the likelihood of individual spots at a bin $g$ can be transformed as follows

$$\begin{aligned}
&\mathbb{P}(\mathbf{X}_{g,I_m}, \mathbf{Y}_{g,I_m}, \mathbf{D}_{g,I_m} \mid \mathbf{Z}_{\cdot,m}, \boldsymbol{\mu}^m, \mathbf{p}^m) \\
=& \prod_{n\in I_m} \mathbb{P}(X_{g,n} \mid Z_{g,m} = k, \mu_k^m)\mathbb{P}(Y_{g,n} \mid Z_{g,m} = k, D_{g,m}, p_k^m) \\
=& \prod_{n\in I_m} \frac{(T_n\mu_k^m)^{X_{g,n}} e^{-T_n\mu_k^m}}{X_{g,n}!} \binom{D_{g,n}}{Y_{g,n}} (p_g^m)^{Y_{g,n}}(1-p_g^m)^{D_{g,n}-Y_{g,n}} \\
=& \Bigg(\prod_{n\in I_m} \frac{T_n^{X_{g,n}}}{X_{g,n}!} \binom{D_{g,n}}{Y_{g,n}}\Bigg)(\mu_k^m)^{\sum_{n\in I_m} X_{g,n}}(p_k^m)^{\sum_{n\in I_m} Y_{g,n}}(1-p_k^m)^{\sum_{n\in I_m} D_{g,n}-Y_{g,n}} \\
=& const \times \frac{(\mu_k^m)^{\sum_{n\in I_m} X_{g,n}} e^{\mu_k^m \sum_{n\in I_m} T_n}}{(\sum_{n\in I_m} X_{g,n})!} \binom{\sum_{n\in I_m} D_{g,n}}{\sum_{n\in I_m} Y_{g,n}} (p_k^m)^{\sum_{n\in I_m} Y_{g,n}}(1-p_k^m)^{\sum_{n\in I_m} D_{g,n}-Y_{g,n}} \\
=& const \times \mathbb{P}(\sum_{n\in I_m} X_{g,n}, \sum_{n\in I_m} Y_{g,n}, \sum_{n\in I_m} D_{g,n} \mid \mathbf{Z}_{\cdot,m}, \boldsymbol{\mu}^m, \mathbf{p}^m).
\end{aligned}$$

Using the equality at each bin $g$, we can rewrite the HMM objective as

$$\begin{aligned}
&\underset{\boldsymbol{\mu}^m, \mathbf{p}^m}{\arg\max} \sum_{\mathbf{Z}_{\cdot,m}} \mathbb{P}(\mathbf{X}_{\cdot,I_m}, \mathbf{Y}_{\cdot,I_m}, \mathbf{D}_{\cdot,I_m} \mid \mathbf{Z}_{\cdot,m}, \boldsymbol{\mu}^m, \mathbf{p}^m))\mathbb{P}(\mathbf{Z}_{\cdot,m}) \\
=& \underset{\boldsymbol{\mu}^m, \mathbf{p}^m}{\arg\max} \sum_{\mathbf{Z}_{\cdot,m}} \prod_{g=1}^{G} \mathbb{P}(\mathbf{X}_{g,I_m}, \mathbf{Y}_{g,I_m}, \mathbf{D}_{g,I_m} \mid \mathbf{Z}_{\cdot,m}, \boldsymbol{\mu}^m, \mathbf{p}^m))\mathbb{P}(\mathbf{Z}_{\cdot,m}) \\
=& \underset{\boldsymbol{\mu}^m, \mathbf{p}^m}{\arg\max}\, const \times \left\{\sum_{\mathbf{Z}_{\cdot,m}} \prod_{g=1}^{G} \mathbb{P}(\sum_{n\in I_m} X_{g,n}, \sum_{n\in I_m} Y_{g,n}, \sum_{n\in I_m} D_{g,n} \mid \mathbf{Z}_{\cdot,m}, \boldsymbol{\mu}^m, \mathbf{p}^m))\mathbb{P}(\mathbf{Z}_{\cdot,m})\right\} \\
=& \underset{\boldsymbol{\mu}^m, \mathbf{p}^m}{\arg\max}\, const \times \left\{\sum_{\mathbf{Z}_{\cdot,m}} \mathbb{P}(\sum_{n\in I_m} \mathbf{X}_{\cdot,n}, \sum_{n\in I_m} \mathbf{Y}_{\cdot,n}, \sum_{n\in I_m} \mathbf{D}_{\cdot,n} \mid \mathbf{Z}_{\cdot,m}, \boldsymbol{\mu}^m, \mathbf{p}^m))\mathbb{P}(\mathbf{Z}_{\cdot,m})\right\}
\end{aligned}$$

□

The above derivation assumes that $\mu_k^n$ in Poisson distribution and $p_k^m$ in Binomial distribution are shared across all spots. But in the mixture of clone cases, the parameters are unique to each spot if the mixing proportions $\theta_{m,n}$ are distinct across spots.

$$X_{g,n} \sim Pois(T_n \sum_m \theta_{m,n}\mu_g^m)$$

$$Y_{g,n} \sim B(D_{g,n}, \sum_m \theta_{m,n} p_g^m, \sum_m \theta_{m,n}(1-p_g^m))$$

But with the given mixing proportion $\theta_{m,n}$, we can pool a subset of spots with similar mixing proportion (and with high tumor proportion) and infer the HMM model only using the pooled subset.

### S10 CalicoST more accurately infers tumor clones and copy number events in simulated data

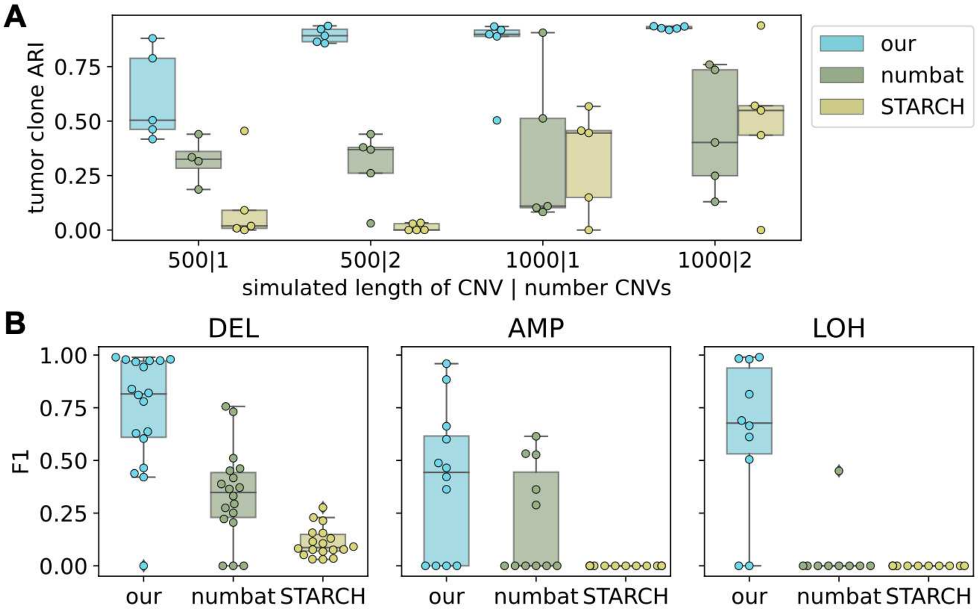


Figure S1: **Accuracy of tumor clone and CNA inference on simulated data.** (A) Accuracy of inferred tumor clones evaluated by ARI of CalicoST, STARCH, and Numbat. X-axis represents simulation settings, including the number of genes spanned by simulated CNA and the number of CNA events per clone, separated by a bar. We simulated multiple samples under each setting. Boxplots and stripplots show the ARI across all simulated samples. (B) F1 score of identifying genes affected by deletions (DEL), amplifications (AMP), and CNLOH events by the three methods. We computed F1 score for each simulated sample under each setting and plotted all samples within each boxplot and stripplot.

It is hard to obtain a ground truth for tumor clone localization in real cancer SRT data, so we use simulated data to evaluate the accuracy of tumor clone identification of CalicoST and compare it with STARCH and Numbat. We simulated three spatially coherent tumor clones and a normal clone within each sample, and each clone contains various settings of simulated CNA events (1 or 2 events, each event spans 500 or 1000 genes). We simulated transcript count matrix from a Dirichlet Multinomial distribution and allele count matrix from a Beta-binomial distribution. We evaluated the accuracy of inferred tumor clone label per spot by adjusted rand index (ARI) compared with the simulated ground truth. We observed that CalicoST overall achieves much higher ARI than STARCH and Numbat across all CNA settings for most of the simulated samples (Fig S1A). Combining the allele frequency (which is absent in STARCH) and spatial information (which is absent in Numbat) improves the tumor clone inference accuracy.

In addition, whether a gene is affected by a CNA event is of interest in SRT data, so we also evaluated the accuracy of identifying genes under deletion (DEL), amplification (AMP), and CNLOH events. We observed that CalicoST achieves a higher F1 score in identifying genes affected by all three types of CNA events (Fig S1B). STARCH does not use allele frequency signals, and thus it cannot identify CNLOH events. Numbat uses allele frequency signals but also has low accuracy in identifying CNLOH. This is potentially explained by the sparsity and few CNAs in the simulation, while CalicoST seeks to reduce the sparsity issue by pooling information across spatially adjacency spots and also across loci in a genomic bin.

#### S11 Running Numbat, STARCH, and InferCNV on HTAN and prostate cancer samples

We ran STARCH, Numbat, and InferCNV on each slide separately to avoid batch effects. For Numbat, we use the default reference expression and prioritize using the recommended running configuration in the "Spatial transcriptomics" tutorial, which sets `max_entropy` to 0.8 and the default parameter for the remaining. But if Numbat reports an error, we modify the `min_LLR` parameter. Table S1 includes the Numbat configurations and final status across slices. For InferCNV, we provide the annotated cell types and specify the normal ones as a reference expression when running on melanoma Slide-tags data, and we specify `analysis_mode = "subclusters"` in order to let InferCNV infer clones. We used the published InferCNV results [44] for the prostate samples in 10X Visium.

Table S1: Running configuration of Numbat

| Slice | Configuration | Status |
|---|---|---|
| CRLM all slices | recommended | Finished |
| prostate H1_2, H1_4 | `min_LLR`=2, `max_entropy`=0.95 | No CNA detected |
| prostate H1_5 | `min_LLR`=2, `max_entropy`=0.95 | Finished |
| prostate H2_1, H2_5 | recommended | Finished |

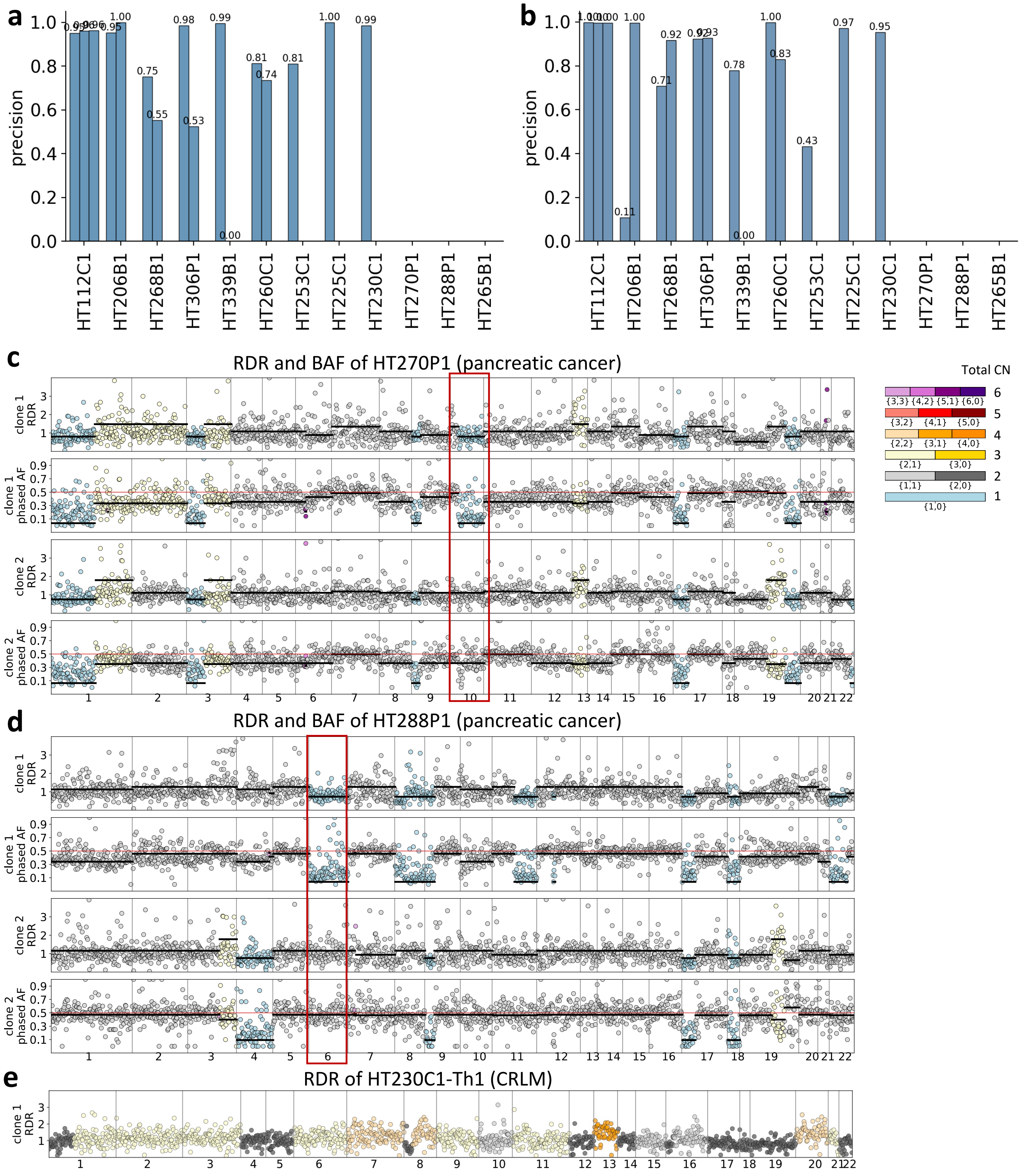


Figure S2: (a–b) Precision (a) and recall (b) of CalicoST-predicted aberrations compared with HATCHet2 identifications from WES data. (c–d) RDR and BAF plots of two pancreatic patients HT270P1 (c) and HT288P1 (d) across CalicoST-inferred cancer clones. Matched WES samples of these two patients have a low tumor purity. Red boxes highlight an example CNA event that distinguishes between clones in each patient. (e) RDR of a CRLM patient HT230C1-Th1 with a triploid genome.

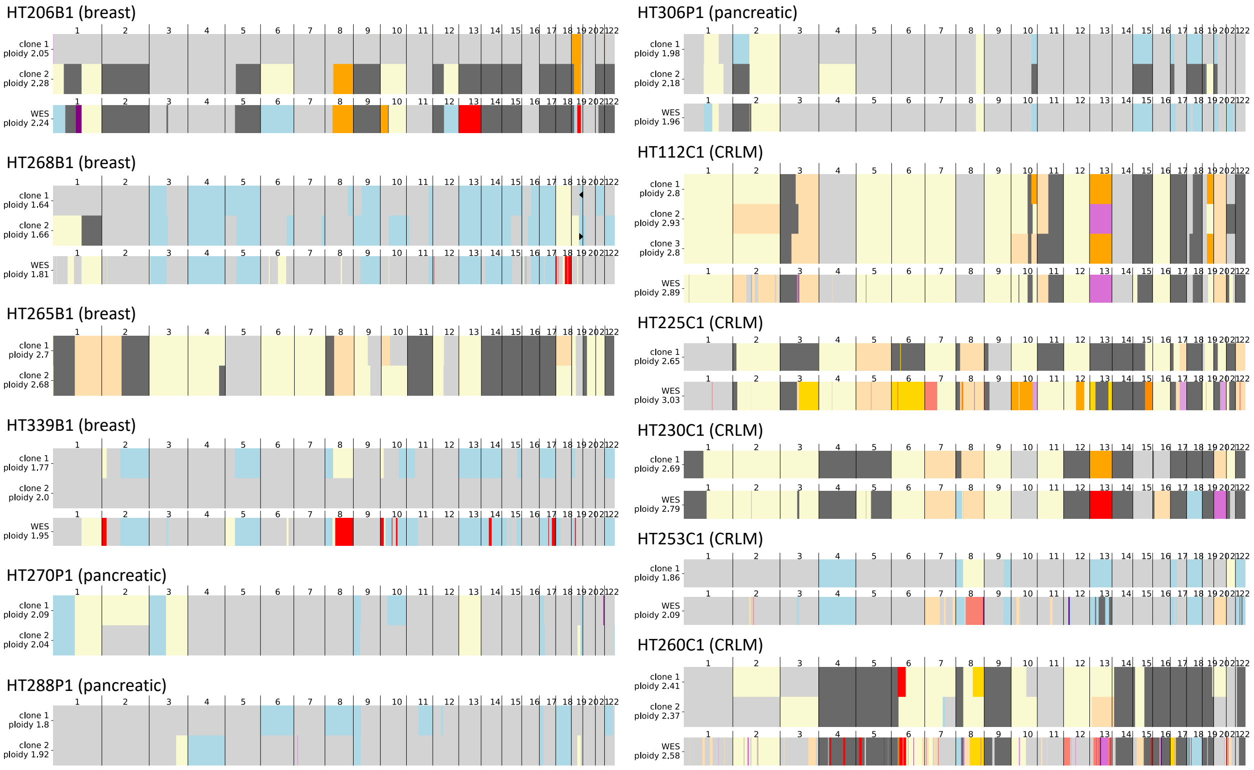


Figure S3: The allele-specific copy numbers inferred by CalicoST from SRT data for twelve patients from HTAN (WashU cohort) and by HATCHet2 from WES data for nine of them.

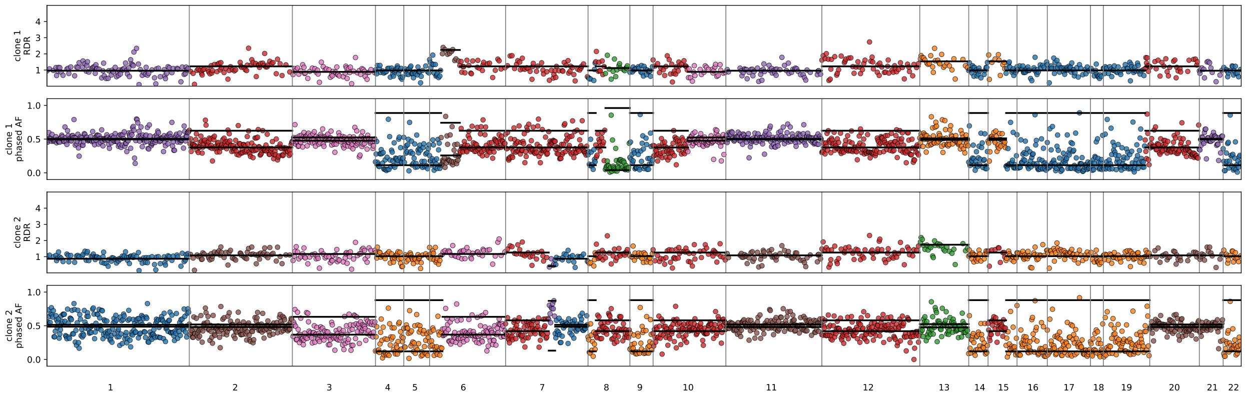


Figure S4: Observed RDR and BAF plots for each inferred clone in sample HT260B1.

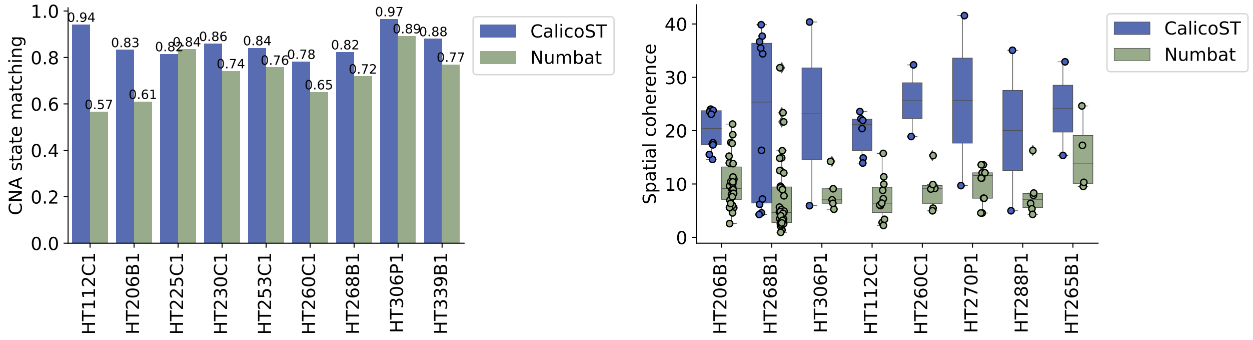


(A) (B)

Figure S5: Accuracy and spatial coherence comparison between CalicoST and Numbat on the all HTAN patients

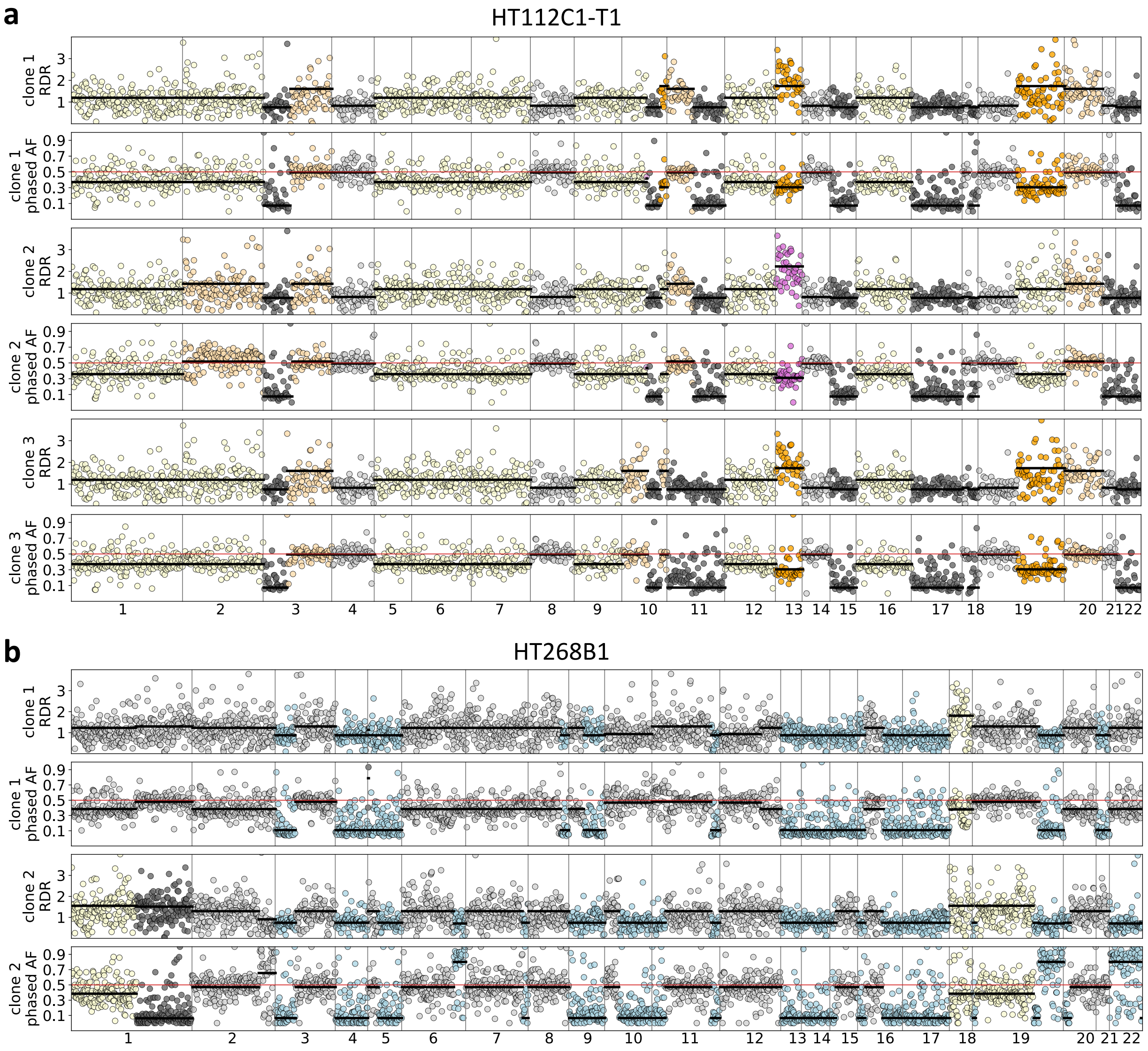


Figure S6: RDR and BAF for each inferred cancer clone of a CRLM patient HT112C1-T1 (a) and a breast cancer patient HT268B1-Th1. (b). X-axis indicates the coordinates along the genome. Color scheme is the same as Fig 4b,d.

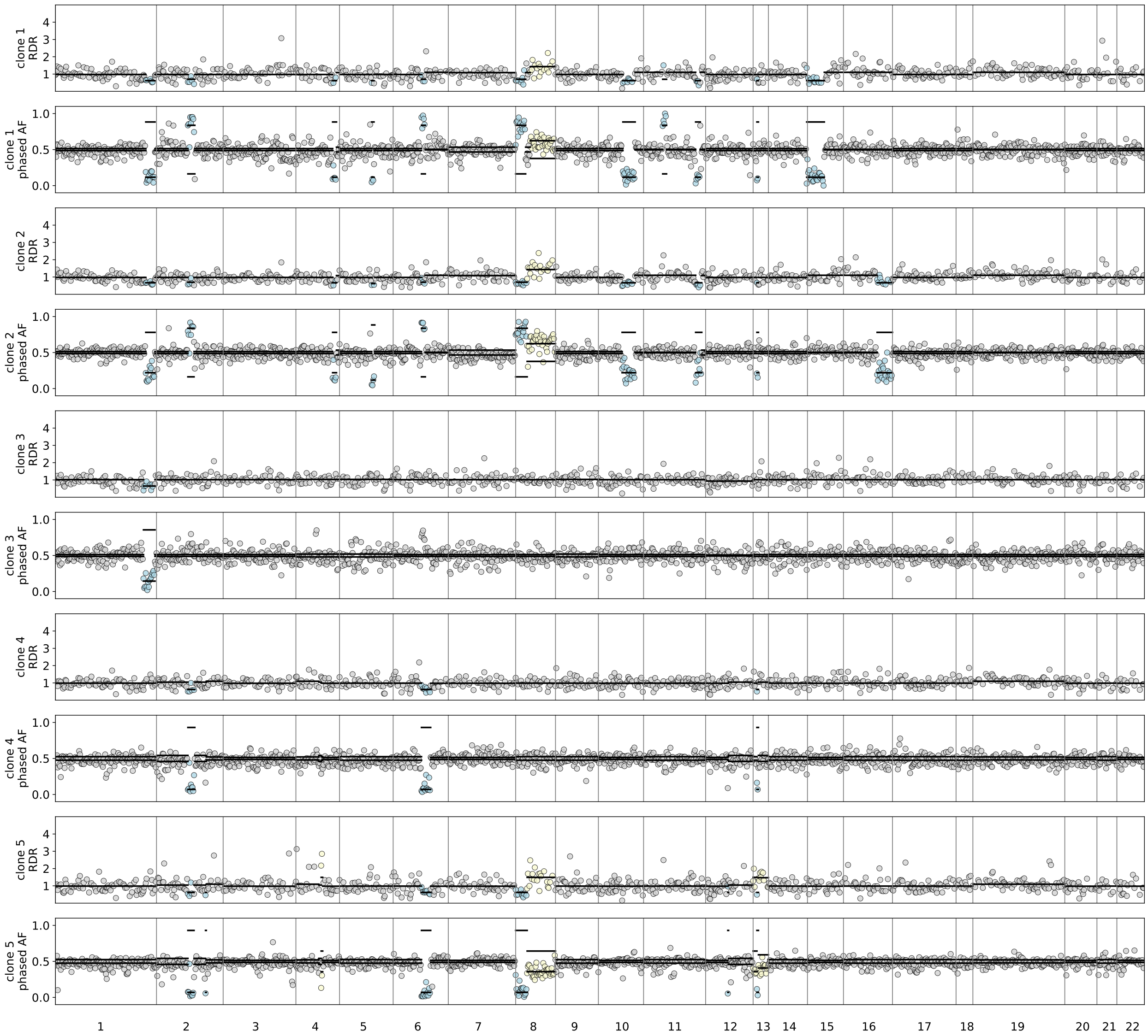


Figure S7: Observed RDR and BAF plots for each inferred clone in the prostate cancer samples.
